# The sensory and motor components of the cortical hierarchy are coupled to the rhythm of the stomach during rest

**DOI:** 10.1101/2021.05.26.445829

**Authors:** Ignacio Rebollo, Catherine Tallon-Baudry

## Abstract

Bodily rhythms appear as novel scaffolding mechanisms orchestrating the spatio-temporal organization of spontaneous brain activity. Here, we follow up on the discovery of the gastric resting-state network (Rebollo et al, 2018), composed of brain regions in which the fMRI signal is phase-synchronized to the slow (0.05 Hz) electrical rhythm of the stomach. Using a larger sample size (n=63 human participants), we further characterize the anatomy and effect sizes of gastric-brain coupling across resting-state networks, a fine grained cortical parcellation, as well as along the main gradients of cortical organization. Most (67%) of the gastric network is included in the somato-motor-auditory (38%) and visual (29%) resting state networks. Gastric brain coupling also occurs in the granular insula and, to a lesser extent, in the piriform cortex. Thus, all sensory and motor cortices corresponding to both exteroceptive and interoceptive modalities are coupled to the gastric rhythm during rest. Conversely, little gastric-brain coupling occurs in cognitive networks and transmodal regions. These results suggest not only that gastric rhythm and sensory-motor processes are likely to interact, but also that gastric-brain coupling might be a mechanism of sensory and motor integration that mostly bypasses cognition, complementing the classical hierarchical organization of the human brain.

**Significance statement:** While there is growing interest for brain-body communication in general and brain-viscera communication in particular, little is known about how the brain interacts with the gastric rhythm, the slow electrical rhythm continuously produced in the stomach. Here, we show in human participants at rest that the gastric network, composed of brain regions synchronized with delays to the gastric rhythm, includes all motor and sensory (vision, audition, touch and interoception, olfaction) regions, but only few of the transmodal regions associated with higher-level cognition. Such results prompt for a reconsideration of the classical view of cortical organization, where the different sensory modalities are considered as relatively independent modules.

## Introduction

Spontaneous brain activity is organized into networks of segregated regions displaying correlated activity across time (Biswal et al., 1995). While the study of resting state networks (RSNs) has proven fundamental to advance our understanding of the functional architecture of the brain (Power et al., 2014), there is still no consensus on RSN exact functions and computations. However, a distinction is often made between networks engaged in sensing and acting on the external world (Yeo et al., 2011), and more cognitive networks that have been associated with a wide spectrum of cognitive processes, ranging from episodic memory, prospective thinking and spontaneous cognition in the default network (Addis et al., 2007; Andrews-Hanna et al., 2014) to saliency detection, cognitive control, and attention in homonymous networks. The distinction between sensory and cognitive networks is not necessarily a sharp one and recently, the spatial layout of RSNs has been reframed in the context of gradients of macroscale cortical organization (Huntenburg et al., 2018) in which most of the spatial variance in cortical functional connectivity (Margulies et al., 2016), myelin (Huntenburg et al., 2017) and gene expression (Krienen et al., 2016) is explained by a connectivity gradient going from default network and transmodal regions to primary sensory-motor cortices.

The division of cortical regions into sensory-motor regions, required for the interaction with the external environment, and transmodal regions, devoted to higher-level cognition, is a recurrent feature not only in the resting-state network literature but also more generally in influential proposals of brain hierarchical organization (Felleman & Van Essen, 1991; Mesulam, 1998; Markov et al., 2013). The organization of the brain in relatively independent sensory and motor modules coordinated by higher-order, transmodal regions leaves little space for the processing of internal bodily information. Despite mounting evidence that bodily rhythms and brain activity at rest are tightly coupled (Azzalini et al., 2019; Tort et al., 2018), RSNs are rarely related to internal, bodily information (for notable exceptions see (Chang et al., 2013; Shokri-Kojori et al., 2018). Recently, we reported the existence of coupling between brain activity at rest and the slow rhythm generated in the stomach (Rebollo et al., 2018; Richter et al., 2017), a finding which was replicated by an independent research group in a single participant scanned multiple times (Choe et al., 2020). The gastric rhythm is a slow (0.05 Hz) electrical oscillation that is intrinsically generated in the stomach wall by a specialized cell type, known as interstitial cells of Cajal (Sanders et al., 2014), and that can be measured non-invasively with cutaneous abdominal electrodes (Koch & Robert M. Stern, 2004; Wolpert et al., 2020). The gastric rhythm is produced at all times but, during digestion, the amplitude of the rhythm increases through modulatory influences of the autonomous nervous system (Cruz et al., 2007; Sveshnikov et al., 2012; Travagli et al., 2006), setting the pace for the contraction of smooth muscles necessary to grind and mix ingested material and eject it into the small intestine.

In the present study, we present a follow-up on the gastric network, a novel resting state network composed of brain regions whose fMRI signals are phase synchronized to the gastric rhythm during resting fixation (Rebollo et al., 2018), in participants who were not in the active phase of digestion. We used a larger sample size (n=63) to provide a detailed characterization in terms of effect sizes and anatomical extent of the human gastric network across the canonical seven resting state networks (Yeo et al., 2011) and across regions of a recent fine-graded multi-modal parcellation of the cerebral cortex (Glasser et al., 2016). We also quantified how the gastric network was positioned along the main two gradients of cortical organization (Margulies et al., 2016). Finally, we explored how different personal, physiological and experimental variables co-varied with the strength of gastric coupling across participants.

## Materials and Methods

### Participants

Seventy-two right-handed human participants took part in this study. Thirty-four participants took part in our first study (Rebollo et al., 2018, sample one) and an additional thirty-eight participants (sample two) were recruited for the present study. All volunteers were interviewed by a physician to ensure the following inclusion criteria: the absence of digestive, psychiatric or neurological disorders; BMI between 18 and 25, and compatibility with MRI recordings. Participants received a monetary reward and provided written informed consent for participation in the experiment and publication of group data. The study was approved by the ethics committee Comité de Protection des Personnes Ile de France III (approval identifier: 2007-A01125-48). All participants were instructed to fast for at least 90 min before the recordings. Data from nine participants were excluded. Three were excluded because excessive head movement during acquisition (translations larger than 3 mm or rotations larger than 3 degrees), three were excluded because their EGG spectrum did not show a clear peak that could allow us to identify the frequency of their gastric rhythm, and three more were excluded because less than 70% of their EGG cycles was within normogastric range (15-30 seconds per cycle, (Wolpert et al., 2020)). A total of 63 participants (mean age 23.95 ± SD 2.76, 31 females, and mean BMI 21 ± SD 1.8) were included in the analysis described below.

### MRI data acquisition

MRI was performed at 3 Tesla using a Siemens MAGNETOM Verio scanner (Siemens, Germany) with a 32-channel phased-array head coil. The resting-state scan lasted 900 s during which participants were instructed to lay still and fixate on a bull’s eye on a gray background. Functional MRI time-series of 450 volumes were acquired with an echo-planar imaging (EPI) sequence and the following acquisition parameters: TR = 2000 ms, TE = 24 ms, flip angle = 78°, FOV = 204 mm, and acquisition matrix = 68×68 × 40 (voxel size = 3×3 × 3 mm^3^). Each volume comprised 40 contiguous axial slices covering the entire brain. High-resolution T1-weighted structural MRI scans of the brain were acquired for anatomic reference after the functional sequence using a 3D gradient-echo sequence. The two samples had different anatomical sequences. The acquisition parameters for the anatomical scan of sample one were the following: TE = 1.99 ms, TR = 5000 ms, TI-1 = 700 ms/TI-2=2500 ms, flip angle-1 = 4°/flip angle-2 = 5°, bandwidth = 240 Hz/pixel, acquisition matrix = 240×256×224, and isometric voxel size = 1.0 mm^3^, total anatomical sequence duration = 11 minutes and 17 seconds. The acquisition parameters of sample two were the following: TE = 3.24 ms, TR = 2300 ms, TI-1 = 900 ms/flip angle-1 = 9°, bandwidth = 210 Hz/pixel, acquisition matrix = 224 × 256×256, and isometric voxel size = 1.0 mm^3^, with a total anatomical sequence duration of 5 minutes and 21 seconds.

### Physiological signal acquisition

Physiological signals were simultaneously recorded during functional MRI acquisition using MRI compatible equipment and the same montage as in Rebollo et al (2018). Briefly, physiological signals were recorded 30 s before and after the resting state acquisition in order to avoid the spreading of the ringing artifact caused by the start of the MRI acquisition. The electrogastrogram (EGG) and electrocardiogram (ECG) were acquired using bipolar EMG electrodes (20 kOhms, 120 cm long) connected to a BrainAmp amplifier (Brain Products, Germany) placed between the legs of participants; the amplifier received a trigger signaling the beginning of each MRI volume. EGG was acquired at a sampling rate of 5000 Hz and a resolution of 0.5 μV/bit with a low-pass filter of 1000 Hz and no high-pass filter (DC recordings). ECG was acquired at a sampling rate of 5000 Hz and a resolution of 10 μV/bit with a low-pass filter of 1000 Hz and a high-pass filter of 0.016 Hz. Prior to the recordings, the skin of participants was rubbed and cleaned with alcohol to remove dead skin, and electrolyte gel was applied to improve the signal-to-noise ratio. The EGG was recorded via four bipolar electrodes placed in three rows over the abdomen, with the negative derivation placed 4 cm to the left of the positive one. The midpoint between the xiphoid process and umbilicus was identified, and the first electrode pair was set 2 cm below this area, with the negative derivation set at the point below the rib cage closest to the left mid-clavicular line. The second electrode pair was set 2 cm above the umbilicus and aligned with the first electrode pair. The positive derivation of the third pair was set in the center of the square formed by electrode pairs one and two. The positive derivation of the fourth electrode pair was centered on the line traversing the xiphoid process and umbilicus at the same level as the third electrode. The ground electrode was placed on the left iliac crest. The ECG was acquired using three bipolar electrodes sharing the same negative derivation, set at the third intercostal space. The positive derivations were set at the fifth intercostal space and separated by 4 cm.

### MRI preprocessing

We used the same MRI preprocessing pipeline as described in Rebollo et al (2018). Brain imaging data were preprocessed using Matlab (Matlab 2017, MathWorks, Inc., United States) and the Statistical Parametric Mapping toolbox (SPM 8, Wellcome Department of Imaging Neuroscience, University College London, U.K.). Images of each participant were corrected for slice timing and motion with six movement parameters (three rotations and three translations). Each participant’s structural image was normalized to Montreal Neurological Institute (MNI) template provided by SPM with affine registration followed by nonlinear transformation (Ashburner et al., 1999; Friston et al., 1995). The normalization parameters determined for the structural volume were then applied to the corresponding functional images. The functional volumes were spatially smoothed with a 3 mm^3^ full-width half-maximum (FWHM) Gaussian kernel.

The time series of voxels inside the brain, as determined using a SPM a priori mask, were subjected to the following preprocessing steps using the FieldTrip toolbox (Oostenveld et al., 2010) (Donders Institute for Brain, Cognition and Behaviour, Radboud University Nijmegen, the Netherlands. See http://www.ru.nl/neuroimaging/fieldtrip, release 06/11/2017). Linear and quadratic trends from each voxel time series were removed by fitting and regressing basis functions, and the BOLD time series were then bandpass filtered between 0.01 and 0.1 Hz using a fourth-order Butterworth infinite impulse response filter. A correction for cerebrospinal fluid motion was obtained by regressing out the time series of a 9 mm diameter sphere located in the fourth ventricle (MNI coordinates of the center of the sphere [0 –46 −32]).

### EGG preprocessing

Data analysis was performed using the FieldTrip toolbox. Data were low-pass filtered below 5 Hz to avoid aliasing and downsampled from 5000 Hz to 10 Hz. In order to identify the EGG peak frequency (0.033–0.066 Hz) of each participant, we first computed the spectral density estimate at each EGG channel over the 900 s of the EGG signal using Welch’s method on 200 s time windows with 150 s overlap. Spectral peak identification was based on the following criteria: peaking power larger than 15μV^2^ and sharpness of the peak. Data from the selected EGG channel were then bandpass filtered to isolate the signal related to gastric basal rhythm (linear phase finite impulse response filter, FIR, designed with Matlab function FIR2, centered at EGG peaking frequency, filter width ±0.015 Hz, filter order of 5). Data were filtered in the forward and backward directions to avoid phase distortions and downsampled to the sampling rate of the BOLD acquisition (0.5 Hz). Filtered data included 30 s before and after the beginning and end of MRI data acquisition to minimize ringing effects.

### Heart rate variability preprocessing

In order to computer the power and ratio of Heart rate variability, we first removed the MRI gradient artefact from the ECG data using the FMRIB plug-in (Iannetti et al., 2005; Niazy, Beckmann, Iannetti, Brady, & Smith, 2005, version 1.21) for EEGLAB (Delorme & Makeig, 2004, version 14.1.1), provided by the University of Oxford Centre for Functional MRI of the Brain (FMRIB). Data from the ECG channels were then bandpass filtered (1–100 Hz) using a FIR filter, designed with Matlab function firws. We then retrieved the inter-beat-interval (IBI) time series by identifying R peaks using a custom semi-automatic algorithm, which combined automatic template matching with manual selection of R peaks for extreme IBIs. Data from eleven participants was discarded at this stage due to noisy ECG recordings. The resulting IBI time series from the remaining 52 participants were then interpolated at 1 Hz using a spline function (order 3), and the average power in the low (0.06-0.15), and high (0.16-0.4) frequency bands was obtained by means of the Fourier transform using Welch’s method on 120 s time windows with 100 s overlap.

### Quantification of gastric-BOLD phase synchrony

We used the procedure described in Rebollo et al (2018) in order to quantify Gastric-BOLD coupling. Briefly, the BOLD signals of all brain voxels were bandpass filtered with the same filter parameters as the ones used for the EGG preprocessing of each participant. The first and last 15 volumes (30 s) were discarded from both the BOLD and EGG time series. The updated duration of the fMRI and EGG signals in which the rest of the analysis was performed was 840 s. The Hilbert transform was applied to the BOLD and EGG time series to derive the instantaneous phases of the signals. The PLV (Lachaux et al., 1999) was computed as the absolute value of the time average difference in the angle between the phases of the EGG and each voxel across time (Equation 1)

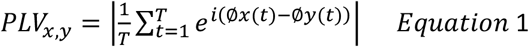

Where T is the number of time samples, and x and y are brain and gastric time-series.

### Statistical procedure for determining regions showing significant gastric-BOLD coupling at the group level

We employed a two-step statistical procedure adapted from previous work (Rebollo et al., 2018; Richter et al., 2017) in order to determine which voxels are significantly coupled to the stomach at the group level. We first estimated chance-level gastric-BOLD coupling at each voxel and in each participant. We then used group-level statistics to determine regions in which gastric-BOLD coupling was greater than chance. In order to estimate the amount of PLV that could be expected by chance, we created a surrogate distribution of PLV using the BOLD signal of one participant and the EGG of the remaining 62 participants. For each participant and voxel, the median value of the surrogate PLV distribution was used as an estimate of chance-level PLV. Since only the empirical PLVs are specific to the frequency and phase of each participant gastric rhythm, our estimate of chance control for biases in PLV that could be expected using physiological signals with the same length, sampling rate, and frequency range as the empirical EGG, but which are not specific to the exact gastric frequency and phase of that participant. Empirical and chance-level PLVs were then compared using a cluster-based statistical procedure that intrinsically corrects for multiple comparisons (Maris and Oostenveld, 2007) implemented in FieldTrip. The procedure consists of applying t-tests between empirical PLV and chance-level PLV across participants at each voxel. Candidate clusters are formed by neighboring voxels exceeding a first-level t-threshold of p<0.01 (two-sided). Each candidate cluster is characterized by the sum of the t-values in the voxels defining the cluster. To determine the sum of t-values that could be obtained by chance, we computed a cluster statistics distribution under the null hypothesis by randomly shuffling the labels ‘empirical’ and ‘chance level’ 1,000 times and applying the clustering procedure. At each permutation, we retained the largest positive and smallest negative summary statistics obtained by chance across all voxels and thus built a distribution of cluster statistics under the null hypothesis, and then assessed the empirical clusters for significance. Because the maximal values across the whole brain are retained to build the distribution under the null hypothesis, this method intrinsically corrects for multiple comparisons. Clusters with a Monte-Carlo p value<0.05 (two-sided, corrected for multiple comparisons) were considered significant and are reported in the Results section.

### Anatomical characterization of the gastric network across resting state networks, cortical gradients, and individual regions

First, we calculated the effect sizes across significant gastric network voxels by computing Cohen’s d on gastric-BOLD coupling strength, defined as the difference between empirical and chance PLV, and then projected the resulting volume in MNI space to Freesurfer‘s average template (Desikan et al., 2006) using registration fusion (Wu et al., 2018). We then used the seven RSN cortical parcellation available in freesurfer native space (Yeo et al., 2011) to identify the networks with more gastric network vertices and with the largest effect sizes. Variability in effect sizes was obtained via bootstrapping. We randomly picked 63 participants with replacement from our original sample, computed Cohen’s d and obtained the standard deviation of this metric across 1000 permutations.

In order to identify the individual regions comprising the gastric network, we used a recent multimodal parcellation of the cerebral cortex (Glasser et al., 2016). This parcellation consists of 180 areas per hemisphere and was obtained using a semi-automatized approach in multi-modal MRI data that detects sharp changes in cortical thickness, myelin, connectivity, and function. The parcellation was imported to freesurfer using the procedure and data available in: https://figshare.com/articles/HCP-MMP1_0_projected_on_fsaverage/3498446/2. For each region of the parcellation, we determined the overlap with the gastric network (percentage of the area in the gastric network), as well as the average effect sizes across significant gastric network vertices.

We then compared the spatial layout of the gastric network with the first two gradients of functional connectivity described in (Margulies et al., 2016), which we retrieved from neurovault (https://identifiers.org/neurovault.collection:1598). We first resampled the volume containing gastric network significant voxels to match the voxel size of the gradients downloaded from Neurovault (3mm^3^ to 2mm^3^), using the tool imcalc from SPM. We then divided each gradient into one hundred equidistant bins and quantified for each bin the percentage of cortical voxels belonging to the gastric network. In order to test whether the gastric network is overrepresented in particular portions of each gradient, we created a distribution (n=1000) of surrogate gastric networks located randomly across the cortex, and recomputed the overlap with each bin and gradient, obtaining a distribution the gastric network across the two gradients under the null hypothesis.

### Variability in gastric coupling across participants and regions

We performed a series of exploratory analyses in order to test for associations between coupling strength and personal (gender, body-mass index, anxiety score), physiological (EGG & heart-rate variability characteristics) or experimental variables (time of day, elapsed time since last meal, head movement, a.k.a frame-wise displacement. Frame wise displacement was estimated as in (Power et al., 2012). State anxiety was measured using the STAI-B questionnaire (Spielberger et al., 1983). Gender and sample effects were assessed using unpaired samples t-test. The influence of the rest of the variables was assessed by separate robust linear regressions using R MASS package (Ripley et al., 2013), which included the covariate and the intercept term. Bayes factor was used to quantify the evidence for H0 relative to H1, with a value <0.33 indicating substantial evidence for a null effect (Kass & Raftery, 2012; Wetzels & Wagenmakers, 2012). For the unpaired samples t-test we used the methods described in (Rouder & Morey, 2011) to determine the Bayes factor. For the regression analysis, an approximation of the Bayes factor was computed (Wagenmakers, 2007) by comparing the Bayesian Information Criterion of the regression model including the covariate and the intercept to a regression model with only the intercept. Group level GLMs on coupling strength across voxels were performed using SPM 12 (Wellcome Department of Imaging Neuroscience, University College London, U.K.). All variables were mean centered and the intercept term was included.

### Code and overlays availability

The code necessary to reproduce the results is available at https://github.com/irebollo/StomachBrain_2021. Unthresholded t-maps, effect sizes and significant voxels can be found at https://identifiers.org/neurovault.collection:9985.

### Visualization tools

The distribution of coupling strength was plotted using raincloud plots (Allen et al., 2019). Visualization of brain data on the cortical surface was done using Pysufer (https://pysurfer.github.io/).

## Results

### The stomach is synchronized with somatosensory, motor, visual and auditory regions

We first determined the frequency of each participant gastric rhythm, by identifying the EGG channel displaying the largest peak within normogastric range (0.033–0.066 Hz). The mean EGG peak frequency across the sixty-three participants was 0.049 Hz (±SD 0.004, range 0.041–0.057). We did not find statistically significant differences between the two samples in EGG frequency (unpaired samples t-test, mean sample one= 0.047 Hz ±SD 0.0034 Hz, mean sample two= 0.048 Hz ±SD 0.0036 Hz, t(61)= 1.41, p=0.161 Bayes factor = 0.6, indicating not enough evidence to disentangle between H1 and H0) nor in EGG power (unpaired samples t-test, mean sample one= 220 µv^2^ ±SD 410 µv^2^, mean sample two= 333 µv^2^ ±SD 530^2^ µv, t(61)= 1.41, p=0.356, Bayes factor = 0.37, indicating not enough evidence to disentangle between H1 and H0). We did find differences in the standard deviation of EGG cycle length, which was significantly smaller in sample two than sample one (unpaired samples t-test, mean sample one= 3.38 seconds ±SD 1.25 seconds, mean sample two= 2.66 seconds ±SD 1.48 seconds, t(61)= -2.08, p=0.041), indicating a more stable gastric rhythm in sample two.

We then quantified the degree of phase synchrony between the EGG signal and BOLD time series filtered around gastric frequency using empirical and chance-level estimates of phase-locking value (PLV). PLV (Lachaux et al., 1999) measures the level of phase synchrony, defined as the stability over time of the time lag between two time series, independently of the amplitude of the two time-series. Importantly, PLV is large as long as the time lag is constant, independently from the value of the time lag, and does not provide information about directionality. In each participant and voxel, we computed the empirical PLV between the narrow-band BOLD signals and the gastric rhythm. We also estimated the amount of PLV that could be expected by chance in each voxel using the BOLD signal of one participant and the EGG of the other sixty-two participants, and taking the median of that surrogate distribution as chance-level PLV. The empirical PLVs were then compared with the chance-level PLVs using a cluster-based statistical procedure that intrinsically corrects for multiple comparisons (Maris & Oostenveld, 2007). Significant phase coupling between the EGG and resting-state BOLD time series occurred in thirty-two clusters (voxel threshold p<0.01, two-sided paired t-test between observed and chance PLV; cluster threshold corrected for multiple comparisons, Monte-Carlo p<0.05). Exact p-values are reported for each cluster in Table S1.

As observed before (Rebollo et al., 2018) the gastric network (Table S1, Figure 1A), comprises bilateral regions along the central, cingulate and lateral sulci, as well as occipito-parietal-temporal regions and portions of the left striatum (Figure 1B), bilateral thalamus (Fig. 1C) and cerebellum (Fig. 1D). The distribution of the average gastric network coupling strength (defined as the difference between empirical and chance PLV) across participants (Fig 1F) ranged from 0.0026 to 0.2096 (mean=0.0417, STD=0.0399, median =0.0319). We found no difference between the two samples in average coupling strength (unpaired sample t-test, mean sample one= 0.043 ±SD 0.044, mean sample two=0.039 ±SD 0.031, t(61)= -0.42, p=0.670, BF=0.27, indicating substantial evidence for H0).

**Figure 1:**
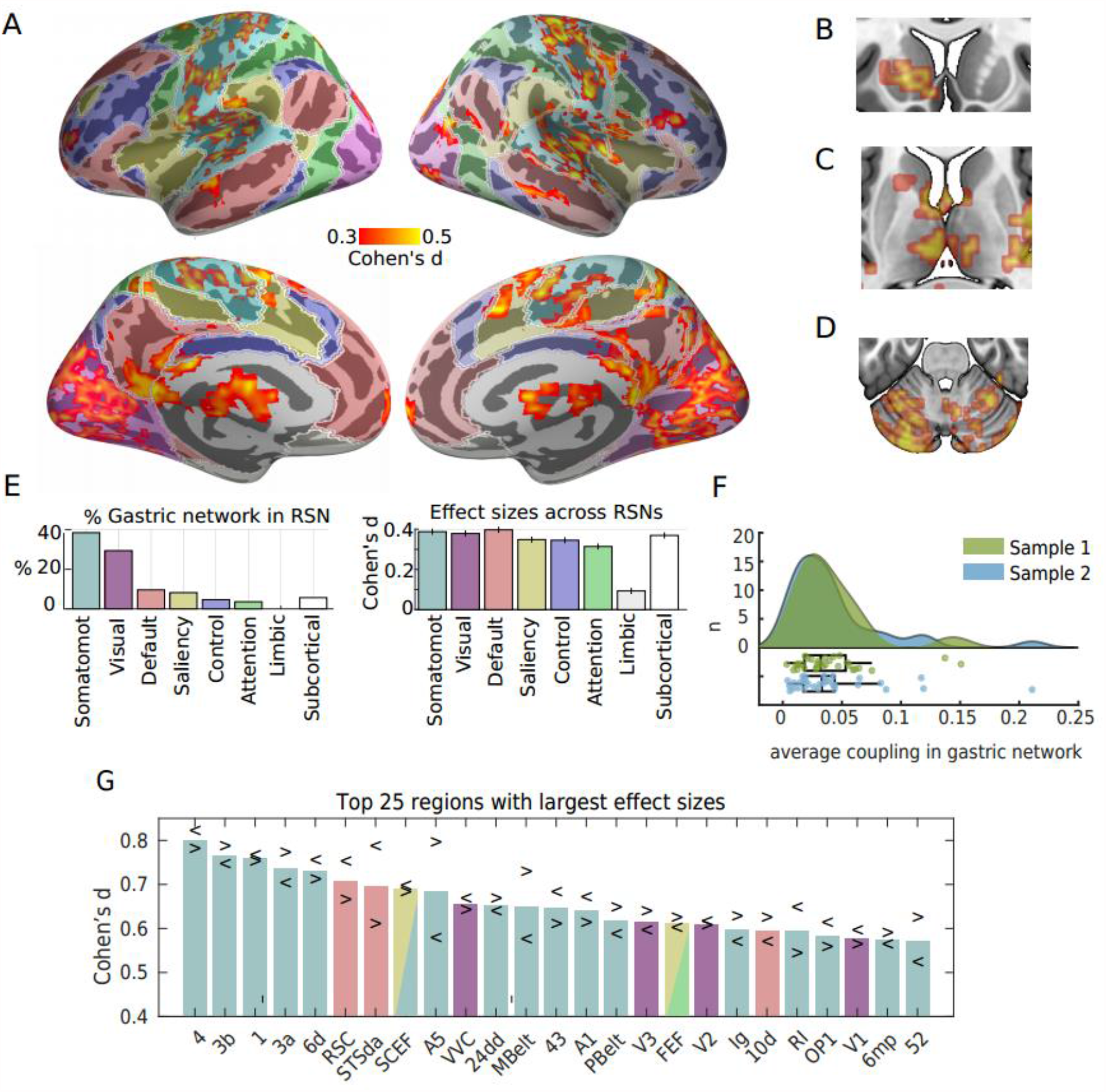
The gastric network and resting state networks. A- Effect sizes (Cohen’s d) of gastric-BOLD coupling are plotted in orange in regions significantly phase synchronized to the gastric rhythm (n=63, voxel-level threshold =0.01 two-sided and cluster significance <0.05 two-sided, intrinsically corrected for multiple comparisons), overlaid on top of the cortical parcellation in seven RSNs proposed by (Yeo et al., 2011), color codes as in E. The gastric network also comprises left striatum (B), bilateral thalamus (C) and cerebellum (D). E- Percentage of the gastric network in each of the brain’s RSNs (left), and average effect size across significant voxels within each network (right). The standard error in effect size was obtained by a bootstrapping procedure. F- Average coupling in the gastric networks across participants, green sample one (n=29), blue sample two (n=34). G- twenty-five regions from (Glasser et al., 2016) parcellation showing the largest effect sizes averaged across hemispheres. Arrowheads depict effect sizes for left (<) and right (>) hemispheres. Abbreviations: RSN, resting state network; Somatomot, somato-motor-auditory.

We then used a well-known parcellation of the cortical surface into seven resting state networks (Yeo et al., 2011) in order to quantify the overlap of the gastric network with canonical resting state networks (Fig 1E). This analysis confirmed that most of the gastric network (67%) is included in the somato-motor (38%) network, which also includes the auditory cortices, and in the visual (29%) network. Gastric coupling in the somato-motor-auditory network spans somatosensory, motor and auditory regions surrounding the central, lateral and cingulate sulci. Gastric coupling in the visual network spans striate and extrastriate visual regions, and is particularly pronounced in medial and ventral occipital regions. Effect sizes (Cohen’s d of paired t-test of empirical against chance PLV) were similar across visual and somato-motor networks (Fig. 1E, right).

The gastric network showed a very limited overlap with non-sensory networks (Fig. 1E, left). The overlap with the default network (9.5%) occurs mostly in one medial node of the default network, the retrosplenial cortex as well as in the lateral node in the superior temporal sulcus, and a small rostral prefrontal region. Only 8.1% of the gastric network is found in the saliency network, sparing core regions of the saliency network such as the anterior insula and the fundus of the dorsal anterior cingulate sulcus. Only very few regions of the gastric network belonged to the control network (4.6%), dorsal attention network (3.6%) or limbic network (0.4%). Subcortical regions (thalamus and striatum), represented 6.8% of the gastric network. Note that although the percentage of the gastric network outside sensory and motor cortices is low, effects sizes in the few coupled regions of the default, saliency, control and attention networks were only slightly smaller than those in sensory and motor cortices (Fig. 1E, right).

To verify that the dominance of sensory and motor cortices in the gastric network was not related to the level of details of the parcellation, we used a more recent parcellation of the cerebral cortex (Glasser et al., 2016) consisting of 180 areas per hemisphere. The parcellation was obtained using a semi-automatized approach in multi-modal MRI data that detects sharp changes in cortical thickness, myelin, connectivity, and function. For each region of the parcellation overlapping with the gastric network, we computed the percentage of the area in the gastric network and the average effect sizes across significant gastric network vertices (Table SII). The 25 regions with the largest effect sizes across both hemispheres (Figure 1 G), were primary motor and somatosensory cortices (Regions 4, 3b,1 and 3a), premotor (6d, SCEF, 6mp, FEF), cingulate motor (24dd), auditory (STSda, A5, MBelt, A1, PBelt, RI, STGa), insular granular, opercular (area 43) early visual regions (V1, V2, V3 and VVC), area FEF from the saliency/attentional networks, and the retrosplenial complex (RSC), area STSda and area 10d, three regions typically associated with the default network.

Because the whole brain analysis indicates that all sensory cortices are coupled to the gastric network except the olfactory cortex, we performed a targeted region of interest analysis of the left and right piriform cortex as defined in the (Glasser et al., 2016) parcellation. We found that coupling in the olfactory cortices has indeed a medium effect size (Cohen’s d left =0.56, right=0.61), comparable to the effect sizes in primary visual (left =0.57, right=0.54) or auditory (left=0.67; right = 0.61) cortices. The absence of olfactory cortices from the whole brain analysis is thus probably due to the small size of those cortices.

### Gastric network and cortical gradients of functional connectivity

The gastric network is mostly found in somato-motor-auditory and visual RSNs. To verify that this is not due to the specific resting-state network parcellations we used, or to a somewhat arbitrary division between sensory and transmodal areas, we analyzed how the gastric network is distributed along the first two gradients of functional connectivity described by (Margulies et al., 2016). In this approach, each cortical voxel can be defined by its location along two different axes, one that goes from unimodal to transmodal regions, and another one that goes from visual to transmodal to somato-motor-auditory regions. In Figure 2A, we reproduce the findings of (Margulies et al., 2016). When considering the whole cortex, the projection of the probability density on the first gradient shows two prominent peaks at the extremities, corresponding to transmodal and unimodal regions (Figure 2A, red curve). When considering only the gastric network, the probability density shows a markedly different profile, with an increase in the unimodal extreme only (Figure 2B, red curve). As described by Margulies et al (2016), the projection of the whole brain probability density on the second gradient shows a prominent peak in transmodal regions (Fig 2A blue curve). However, gastric network voxels are more densely represented in the visual and somato-motor-auditory extremes of the gradient (Fig 2B blue curve), indicating that coupling with the gastric rhythm is more likely to be present in unimodal than in transmodal brain regions.

**Figure 2:**
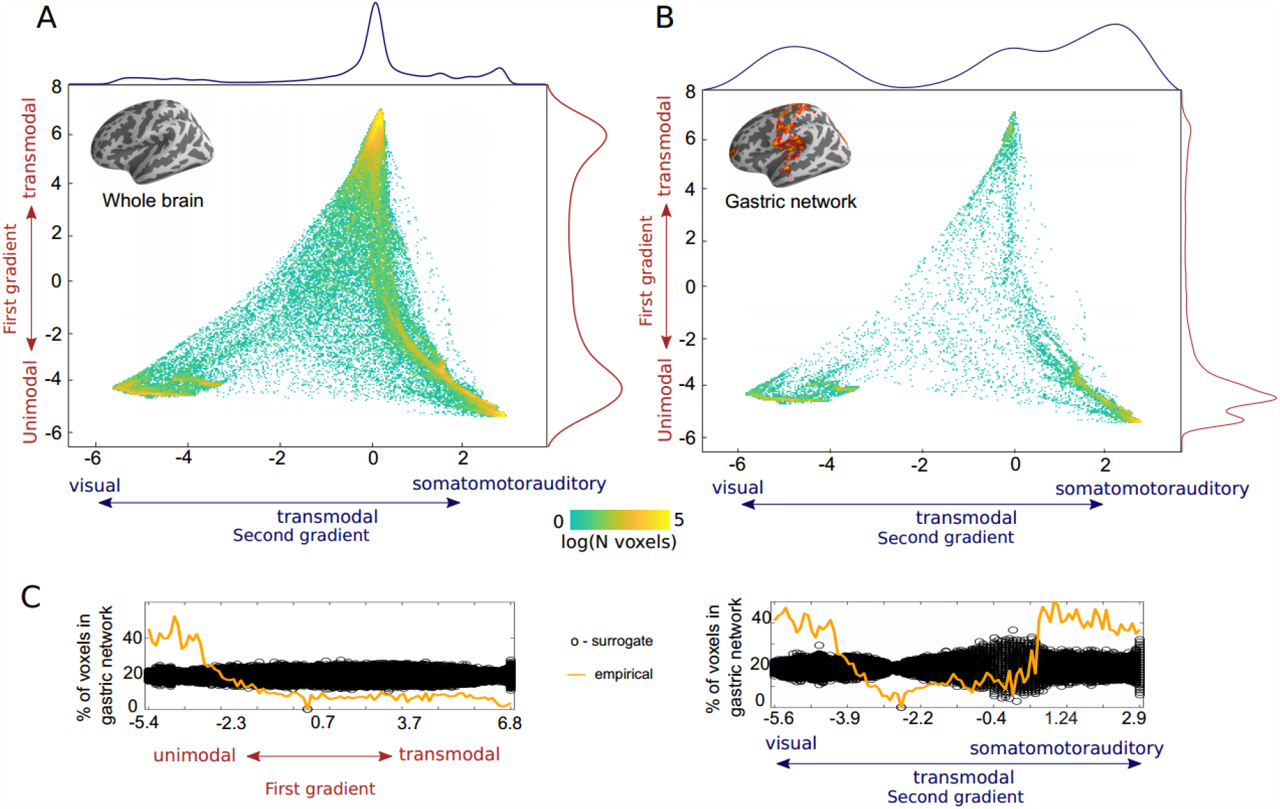
The gastric network and cortical gradients of functional connectivity. A- Density plot depicting the distribution of all cortical voxels along the first two gradients of functional connectivity described by (Margulies et al., 2016). The first gradient (y-axis), runs from unimodal (negative values, sensory or motor regions) to transmodal regions (positive values). The second gradient (x-axis) runs from visual (positive values) to somato-motor and auditory regions (negative values). The color scale depicts the logarithm of the number of voxels. The projection of the probability density on the first and second gradients are shown in red and blue respectively. B- Density plot of gastric network voxels on the first two gradients of functional connectivity. Gastric network voxels are located in the unimodal extremes of the two gradients. The projection of the probability density on the first and second gradients are shown in red and blue respectively. C- Percentage of all brain voxels that belong to the gastric network (orange) for each of the two gradients, computed on one hundred equidistant bins. The black circles depict the distribution of chance level overlap obtained by reallocating randomly the spatial position of gastric network voxels in the cortex. Gradients downloaded from https://identifiers.org/neurovault.collection:1598

We then tested whether the distribution of gastric network voxels along the first and second gradients could be due to chance. We first computed the percentage of brain voxels that belonging to the gastric network for each of the two gradients across one hundred equidistant bins (Fig 2C, orange lines). In order to test whether spatial biases of such size could be obtained by chance, the spatial position of the gastric network voxels was randomly permuted across the cortex, while keeping the spatial layout of the gradients intact, and computed the percentage of voxels belonging to the surrogate gastric network thus created. The procedure was repeated one thousand times to estimate the distribution under the null hypothesis (Fig 2C, black circles). For the first gradient, the percentage of all brain voxels overlapping with the gastric network across bins is systematically larger than chance in the unimodal bins of the gradient, and systematically smaller in the remaining bins (Fig 2C left). Similarly, for the second gradient, the overlap with the gastric network across bins is systematically larger than chance in the unimodal bins and systematically smaller in the transmodal bins (Fig 2C right). This analysis confirmed that the spatial biases in the unimodal extremes of the gradients could not be obtained by chance.

### Anatomical characterization and effect sizes in the gastric network

We now examine in more detail the anatomy of the gastric network, starting with regions belonging to the somato-motor network in central and mid-cingulate regions, followed by a characterization of opercular regions in and around the lateral sulci, including auditory cortices and insula, then regions located in the posterior portion of the medial wall and in occipital, posterior cingulate, temporal and parietal cortices, and ending with the few transmodal regions in frontal, prefrontal and lateral temporal cortices.

Gastric network in central and mid-cingulate regions In central and mid-cingulate regions (Figure 3A), the gastric network covers primary somatosensory (green), motor, premotor, and cingulate motor (blue) and non-motor (violet) cingulate regions, as well as area 55b (pink). The primary somatosensory cortex proper (areas 3b and 1), is located in the postcentral gyrus. Accessory somatosensory area 2, in the bank of the post-central sulcus, shows little coupling with the gastric rhythm. In the anterior direction, area 3a, in the fundus of the central sulcus, is sometimes referred to as part of the somatosensory cortex but can also be seen as a transition zone with primary motor area 4 (Catani, 2017) in the posterior bank of the postcentral gyrus. Both area 3a and area 4 display coupling with the gastric rhythm. The gastric network is also found in numerous premotor areas: area 6 (6m, 6d, 6v, 6a), the frontal eye field (FEF) and supplementary motor areas (6mp, 6ma, SCEF). Note that area SCEF (supplementary motor and cingulate eye fields), extends into the dorsal bank of the anterior cingulate sulcus. Along the medial wall, the gastric network overlaps with left area 5l (cyan), the cingulate motor areas (24dd, 24dv, blue), the paralimbic cortex (area p32pr), areas p24pr and 23d, located adjacent to the corpus callosum and area 23c, located between the cingulate motor regions and the task-negative precuneus Finally, the gastric network also includes area 55b mostly on the right side, a recently discovered region which is left-lateralized for language and right-lateralized for theory of mind (Glasser et al., 2016), and to a lesser extent, the superior frontal language area (SFL), a region that is also left-lateralized for language.

**Figure 3:**
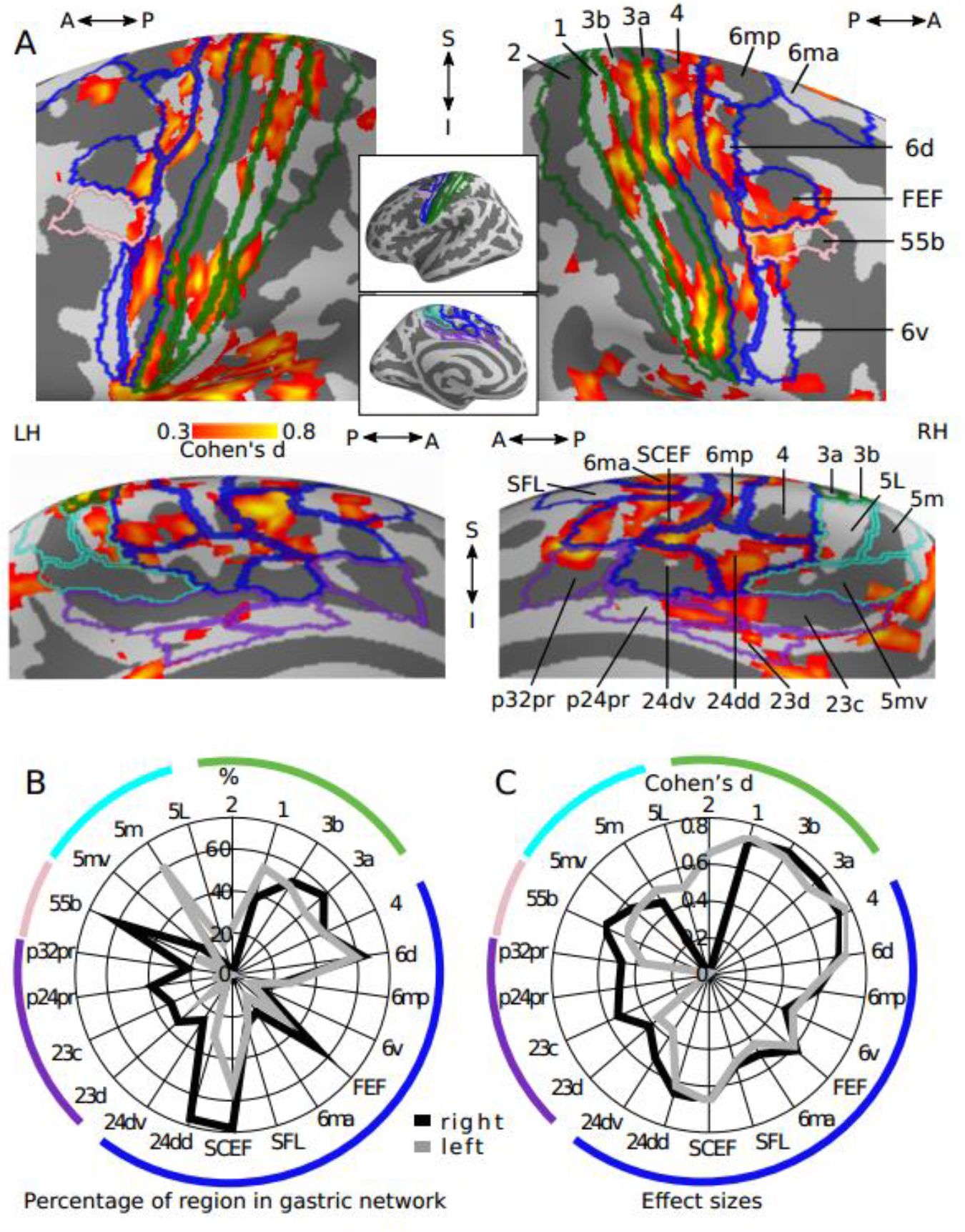
Gastric network in central and mid-cingulate regions. A- Effect sizes in the gastric network in central (top) and mid-cingulate regions (bottom) displayed on the left and right inflated surfaces and overlaid with the corresponding regions of Glasser et al 2016 parcellation. Green, primary somatosensory; Blue, motor and premotor regions; Pink, area 55b; Violet, non-motor cingulate regions; Cyan, area 5 and its subdivisions. B- Percentage of each region overlapping with the gastric network. C- Effect sizes of gastric-BOLD coupling in voxels belonging to the gastric network split by regions. Abbreviations as in text and table SII.

Regions containing a large proportion of voxels belonging to the gastric network (Figure 3B) include primary somatosensory (3b, 3a, 1), primary motor (4), premotor (SCEF, 6d, FEF), cingulate motor regions (24dd), region 55b and area 5m. Gastric-brain coupling show a strong right lateralization in regions surrounding the cingulate sulcus (p32pr, p24pr, 23cc 23d, 24dv, 24dd, SCEF), as well as in FEF and area 55b, while areas 1 and 5m are left-lateralized. Effect sizes (Figure 3C) were largest in somatosensory (3b, 1, 3a) and primary motor (4) cortices, followed by premotor and cingulate motor regions (6d, SCEF, 24dd, FEF, 6m). Asymmetries in effect sizes showed similarities with asymmetries in overlap, with right lateralization in effect size in cingulate regions (p24pr, 23cc 23d, 24dv, 24dd, SCEF) and region 55b, and left lateralization in regions 5l, 5m and 2.

### Gastric network surrounding the lateral sulcus

In and around the lateral sulcus (figure 4A), the gastric network spans secondary somatosensory (green), frontal operculum (blue), early (dark violet) and association (light violet) auditory regions as well as regions of the insula proper (pink). The secondary somatosensory (SII) is composed of areas OP1, OP2-3, and OP4, and located anteriorly to area Pfcm, in the inferior parietal cortex. SII is separated from the frontal operculum (areas FOP1, FOP2, FOP3, and FOP4) by area 43 (cyan). Early auditory regions include the primary auditory cortex (A1) and the surrounding lateral belt (LBelt), medial belt (MBelt), parabelt (PBelt) and retro-insular cortex (RI). Auditory association regions extend inferiorly to the superior temporal sulcus and include A4, A5, Temporal Area 2 (TA2), and auditory default network regions dorsal posterior superior temporal sulcus (STSdp), dorsal anterior superior temporal sulcus (STSda), and ventral anterior superior temporal sulcus (STSva). The insula proper consists of area 52, a transition region between auditory cortex and insula, the granular insula (Ig), which contains a complete somatotopic motor map (Glasser et al., 2016), the middle insular area (MI), and posterior insular area I and II (PoI1, PoI2).

**Figure 4:**
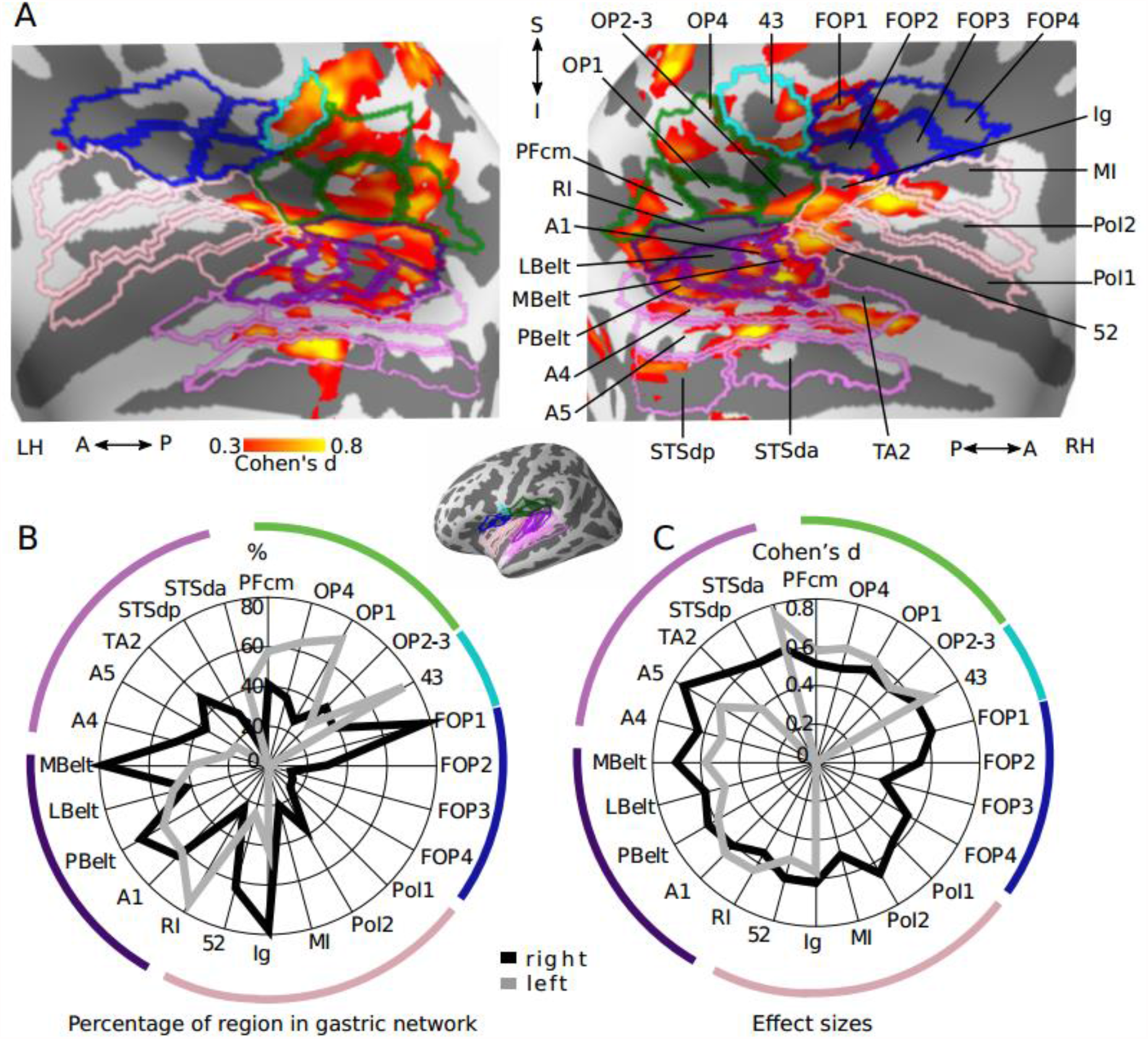
Gastric network in and around lateral sulcus. A- Effect sizes in the gastric network in and around left and right lateral sulci, displayed on inflated cortical surfaces and overlaid with the corresponding regions of the Glasser 2016 parcellation. Green, secondary somatosensory cortex and area PFcm; Blue, frontal operculum; Cyan, area 43; Dark violet, early auditory; Light Violet, auditory association; Pink, Insula proper. B- Percentage of each region overlapping with the gastric network. C- Effect sizes of gastric-BOLD coupling in voxels belonging to the gastric network split by regions. Abbreviations as in text and table SII.

The regions with the largest overlap with the gastric network consist of early auditory regions (Pbelt, A1, MBelt, RI), right insular regions (Ig, 52), left parietal opercular regions (OP4, OP1, 43) and right area FOP1 (Figure 4B). Effect sizes were largest in early and association auditory regions (A5, A4, MBelt, LBelt, A1, PBelt, STSda, RI, TA2), insular cortex (52, Ig, PoI2) and areas 43 and FOP1 (Figure 4C). Overall, overlap and effect sizes in SII (OP1, OP2-3) were most prominent in the left hemisphere while overlap and effect sizes in the frontal, insular, MBelt and auditory association regions were strongly right-lateralized.

### Gastric network in posterior regions

In the posterior part of the brain, the gastric network covers large portions of the occipital visual cortex, (Figure 5A, dark green), occipito-parietal sulcus (blue) and retrosplenial cortex (violet), dorsal precuneus (light green), ventral visual stream (pink), and right temporo-parietal-occipital junction (black). The gastric network is found in all early visual regions (V1, V2, V3, and V4), extending superiorly to the dorsal stream (V3A, V3B, V6, V6A, V7), up to area 7Am and the precuneus visual area (PCV). Medially and anteriorly, the gastric network extends to the parieto-occipital sulcus and recruits portions of the dorsal visual transition area (DVT), prostriate cortex (Pros), and parieto-occipital sulcus regions 1 and 2 (POS1, POS2). Area POS2 stands out as due to its large overlap with the gastric network (90%). Conversely, it is worth noting that the gastric network spares most of the ventral precuneus (Yellow; areas 7m, 31pd, 31a, 31pv), which is one of the core nodes of the default network. Ventrally, the gastric network extends to ventromedial visual areas 1, 2 and 3 (VMV1, VMV2, and VMV3), as well as regions of the ventral stream (V8, area lateral occipital 2, LO2, posterior inferotemporal complex, PIT, and fusiform face complex, FFC), and the occipital-temporal-parietal junction (Black, MT, Pgp, TOPJ3, LO3) in the right hemisphere.

**Figure 5:**
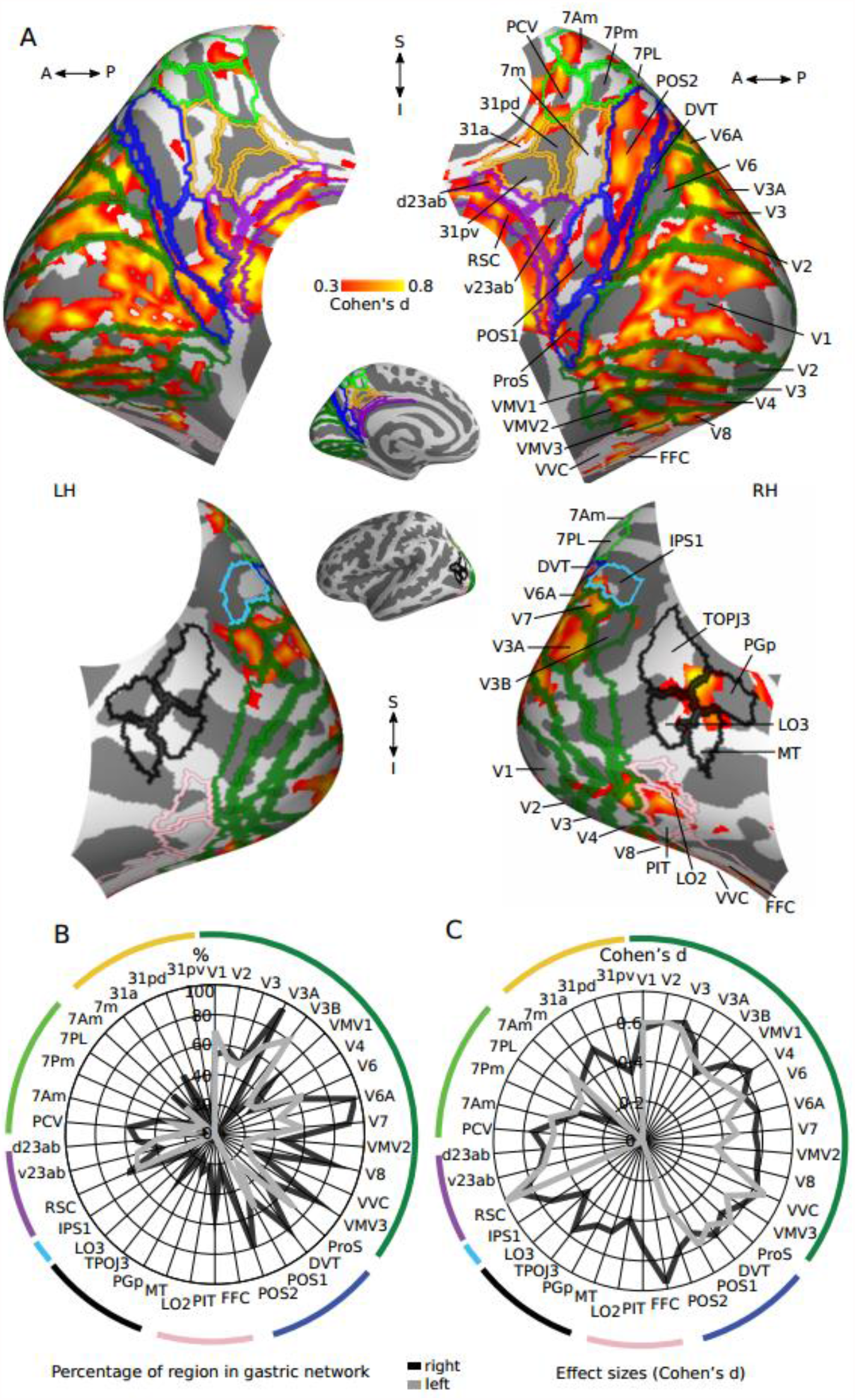
Gastric network in posterior regions. A- Medial (top) and lateral (bottom) views of effect sizes in the gastric network in left and right posterior regions, displayed on inflated cortical surfaces and overlaid with the corresponding regions of Glasser 2016 parcellation. Dark green, early visual; Blue, occipito-parietal sulcus; Violet, retrosplenial and posterior cingulate cortices; Yellow, ventral precuneus; Light Green, dorsal precuneus; Black, temporo-parietal-occipital junction; Pink, lateral occipital and fusiform. B- Percentage of voxels belonging to the gastric network, in each region. C- Effect sizes of gastric-BOLD coupling in voxels belonging to the gastric network split by regions. Abbreviations as in text and table SII.

A cluster of dorsal right regions (V3A, V6A and V7, POS2 and DVT) stands out, with more than 80% of those regions in the gastric network (Figure 5B). Other posterior regions with large overlap were right area VMV3, left V3B, and, to a lesser extent, the RSC, as well as left V1 and ProS. Effect sizes (Figure 5C) were largest in RSC, right FFC and most occipital visual regions including V1. A rightwards lateralization in effect sizes and overlap was present for most regions, with the exception of area V3B, which displayed a leftwards lateralization.

### Gastric network in prefrontal and lateral temporal cortex

The gastric network includes a few transmodal regions located in the middle temporal gyrus (Figure 6A, pink), inferior frontal cortex (blue), dorsolateral prefrontal cortex (green), and in the orbitofrontal cortex (yellow). In the middle temporal gyrus, the gastric network is found in the middle temporal gyrus (TE1m, TE1p), as well as in ventral regions of the superior temporal sulcus (STSdp, STSvp), belonging to the default network. The percentage of voxels belonging to those regions remain small (Figure 6B), with moderate effect sizes (Figure 6C), and a marked right lateralization.

**Figure 6:**
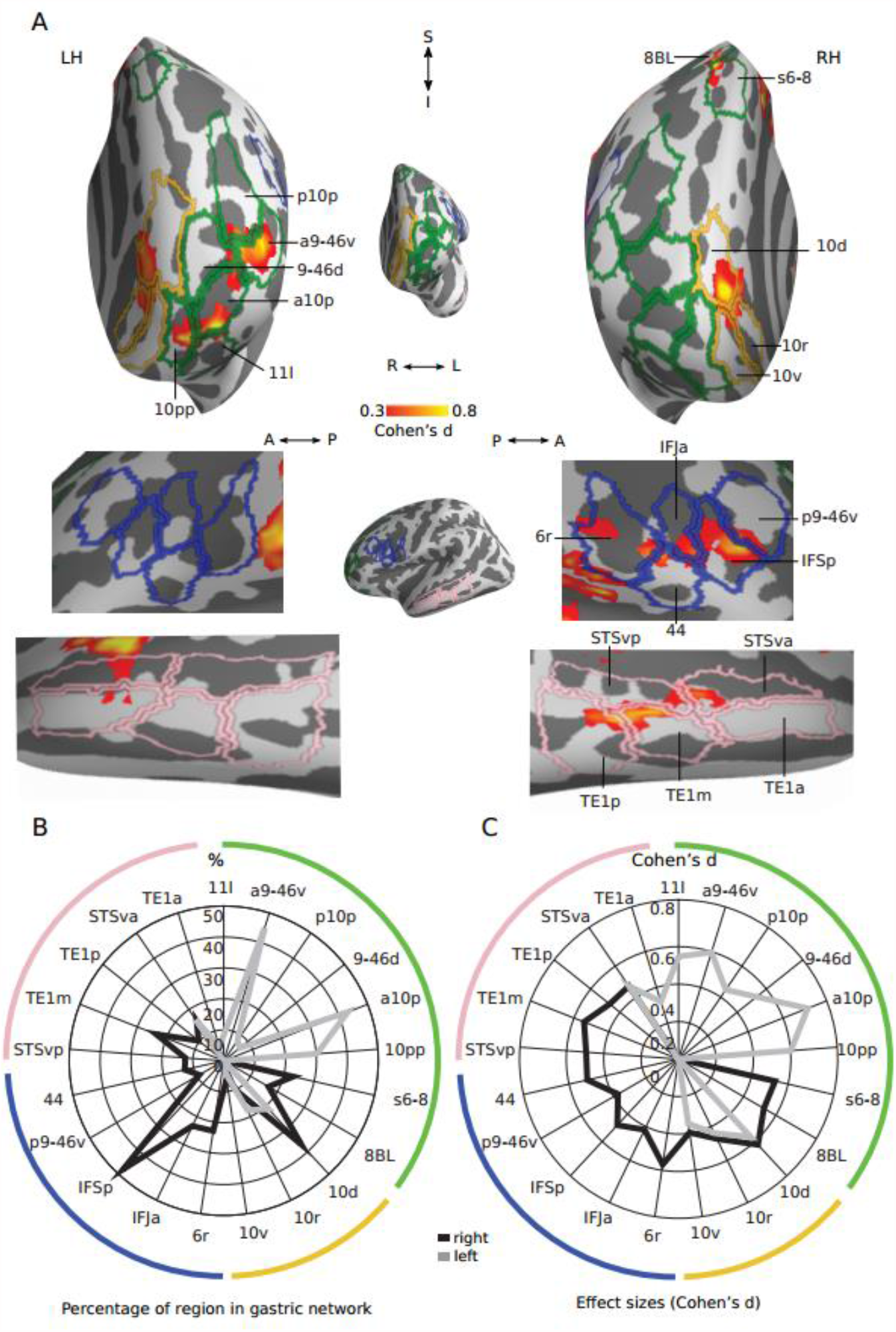
Gastric network in prefrontal and lateral temporal. A- Frontal (top) and lateral (bottom) views of effect sizes in the gastric network along left and right prefrontal and lateral temporal regions, displayed in inflated cortical surfaces along with the corresponding regions of Glasser 2016 parcellation. Light Green, lateral prefrontal cortex; Yellow, orbitofrontal cortex; Blue, inferior frontal gyrus; Pink, superior temporal sulcus. B- Percentage of each region belonging to the gastric network. C- Effect sizes of gastric-BOLD coupling in voxels belonging to the gastric network split by regions. Abbreviations as in text and table SII.

In right inferior frontal cortex (Figure 6A, blue), the gastric network forms a cluster that comprises the posterior portion of the inferior frontal sulcus (IFSp), the anterior portion of the inferior frontal junction (IFJa), and area posterior 9-46d (p9-46d) as well as area 44, all regions of the control network, extending to rostral area 6 (6r) of the saliency network. In the left dorsolateral prefrontal cortex (Figure 6A green), the gastric network includes multiple regions of the control network, including Superior Transitional Area 6-8 anterior (s6-8a), anterior area 10p (a10p), area 11 lateral (11l), anterior area 9-46 ventral (a9-46v), area 9-46 dorsal (9-46d), as well as area 8b lateral from the default network and polar area 10p (p10p) from the limbic network. Overlap is particularly pronounced in a10p and a9-46v (Figure 6B) with an exclusively left lateralization. Finally, in orbitofrontal cortex (Figure 6A yellow), the gastric network includes dorsal and rostral portions of area 10 (10d, 10r) from the default.

### Variability in gastric coupling across participants and regions

We examined how differences in personal (gender, body-mass index, anxiety score), physiological (EGG & heart-rate variability characteristics) or experimental variables (time of day, elapsed time since last meal, head movement, a.k.a frame-wise displacement), were related to variability in coupling strength across participants, in a series of exploratory analyses. Due to the small age range of our sample (18-30), we did not include age as a regressor.

We first tested whether any of these variables accounted for a compact measure, the mean coupling strength in the gastric network (Table 1). We found no differences between genders in the average coupling strength (mean female= 0.039±SD 0.036, mean male = 0.044 ±SD 0.044, paired t-test, t(61)= 0.48, p=0.634, Bayes factor = 0.279, indicating substantial evidence for the null hypothesis). All other variables were tested with robust linear regressions. With one exception, none of the variables tested accounted for coupling strength, including state anxiety (t(58)= 0.47, p=0.634, Bayes factor=0.124 indicating substantial evidence for the null hypothesis). The exception was the ratio of low over high-frequency heart-rate variability (LF-HRV/HF-HRV, Figure 7C), which is associated with larger brain coupling (n=52, t(50)= 3.25, R= 0.2, uncorrected p= 0.002, Bonferroni corrected p= 0.026). Note that LF-HRV/HF-HRV indexes the relative contribution of different autonomic control mechanisms on heart rate, but provides no clear-cut distinction between sympathetic and parasympathetic activity (Goldstein et al., 2011).

**Table 1:**
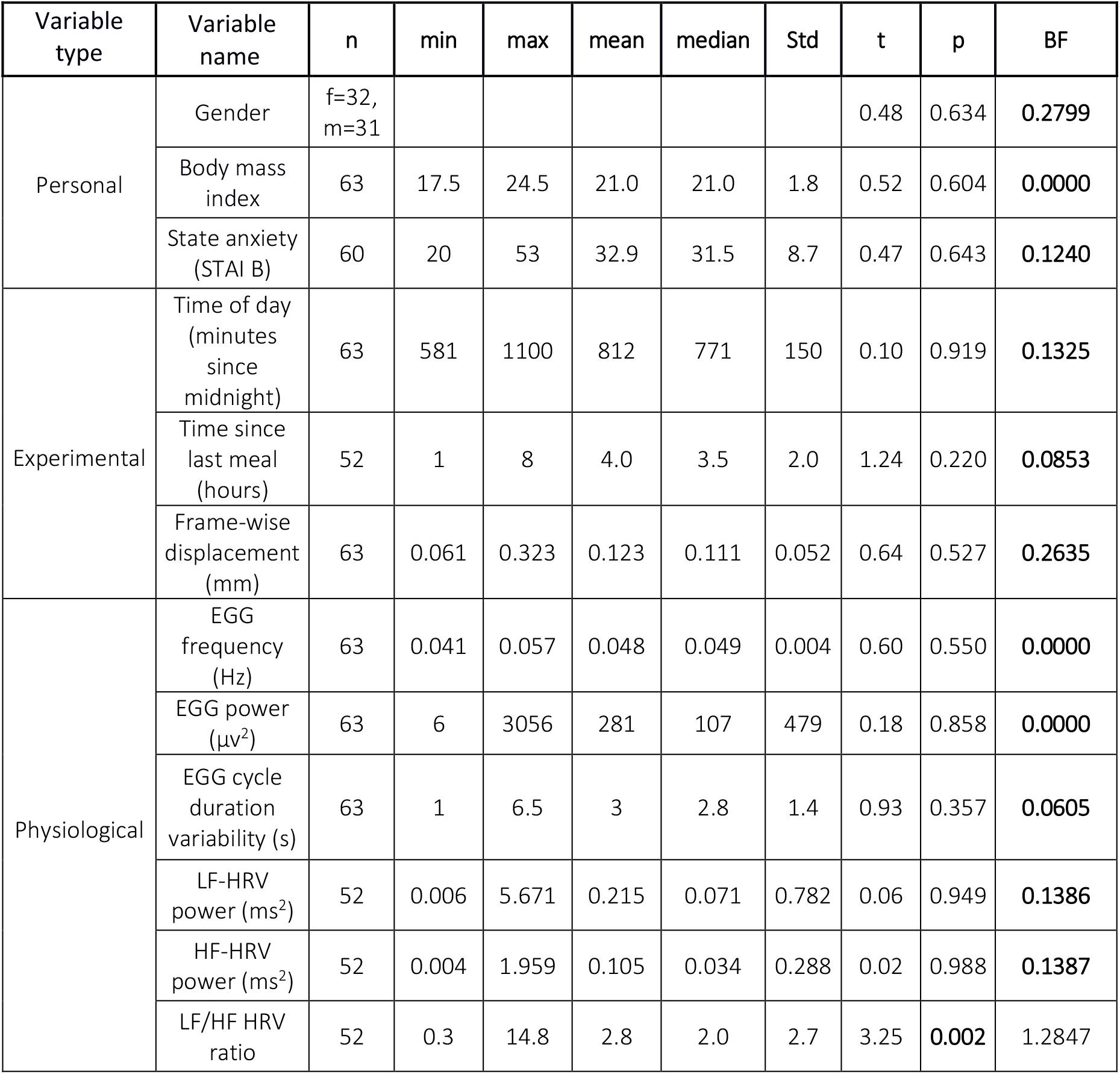
Variables tested to account for average coupling strength in the gastric network. For each variable tested the table displays the descriptive statistics, t-values, uncorrected p values, and Bayes factors (BF), for which values smaller than 0.33 indicates substantial evidence for the null hypothesis. Gender effects were assessed by comparing males and females with an unpaired sample t-test, all other variables were assessed using robust linear regressions. Bold font indicates significant differences from the null distribution for p values and substantial evidence for the null hypothesis for Bayes factors.

**Figure 7.**
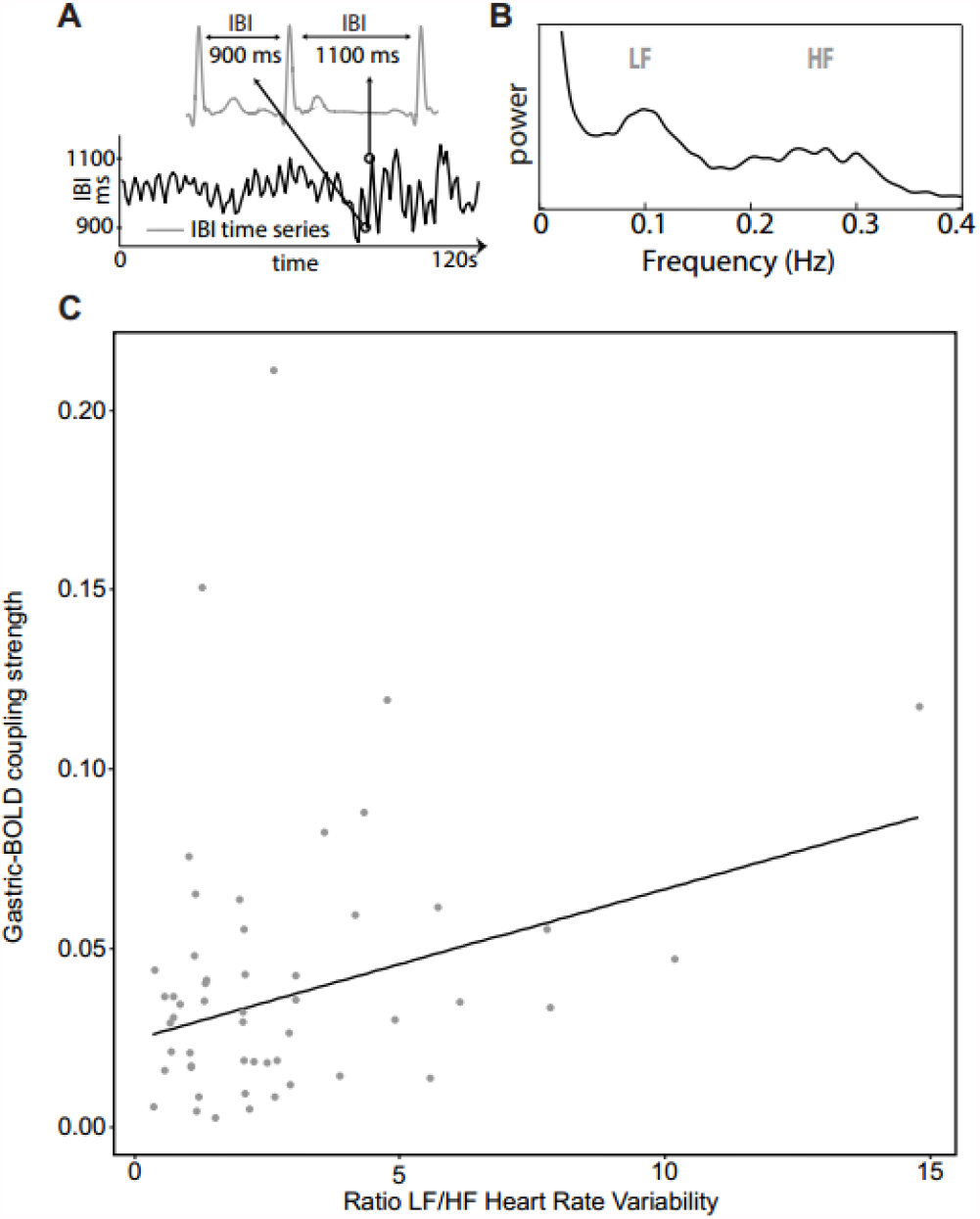
Association between coupling strength and autonomic activity. A- Inter-beat intervals (IBI), or time difference between each heart-beat, are used to build time series of heart rate variability. Reproduced from (Azzalini et al., 2019). B. The power spectrum of heart-rate variability, displaying a prominent peak in the low-frequency band (LF, 0.05-0.15 Hz), reflecting both sympathetic and parasympathetic activity, and in the high-frequency band, (HF, 0.16-0.25 Hz), reflecting parasympathetic activity. C- A robust linear regression shows a significant slope between the average coupling strength in the gastric network and the LF-HRV/HF-HRV.

We reasoned that only some gastric network regions might be modulated by the variables we analyzed. We therefore used two different strategies to look for more subtle effects of inter-individual variables on coupling strength, both accounting for multiple comparisons. We first tested for effects of these variables across Glasser et al (2016) ROIs overlapping with the gastric network, applying false-discovery rate correction over the 226 ROIs tested for each variable separately. We only found one significant association, with coupling strength in right area 7Am co-varying with EGG cycle duration variability (t(61)=3.936, FDR p = 0.048). We also performed group level general linear models with coupling strength in all brain voxels as the dependent variable. Gender, group, BMI, frame-wise displacement, time of the day, EGG frequency, EGG power were used in a first general linear model with all participants (n=63). Separate general linear models including these variables and either state anxiety (n=61), elapsed time since last meal (n=51) or heart rate variability measures (n=51) were performed. None of the models survived correction for multiple comparisons (FWE p<0.05).

## Discussion

We examined the anatomy of the gastric network, a set of brain regions phase coupled to the rhythm of the stomach during rest, and analyzed the spatial layout of the gastric network at the level of its constituent regions (Glasser et al., 2016), resting-state networks (Yeo et al., 2011), and its extent and position along the first two gradients of cortical connectivity that underlie the topological structure of the cortex (Margulies et al., 2016). We found that the gastric network is overrepresented in unimodal sensory-motor regions, and underrepresented in transmodal regions. All sensory and motor cortices are coupled to the gastric rhythm, including not only areas responding to touch, vision and audition but also the interoceptive insula and, to a lesser extent, the olfactory piriform cortex. Only few transmodal regions were coupled to the gastric rhythm, mostly in the default network (retrosplenial cortex, right area STSda). None of the personal, physiological and experimental variables tested co-varied with the overall strength of gastric coupling across participants, with the exception of an index of cardiac autonomic activity. Notably, we found substantial evidence for an absence of association between anxiety and gastric-brain coupling during rest.

### Anatomical pathways for gastric brain coupling

Some of the gastric network regions we observe are known to be involved in interoception and autonomic functions. Such is the case of the granular insula, the somatosensory cortices, and the cingulate motor regions (Amassian, 1951; Cechetto & Saper, 1987; Dum et al., 2009), which receive visceral inputs, and motor regions, which provide sympathetic input to the stomach (Levinthal & Strick, 2020). However, we find that gastric-brain coupling extends well beyond expected visceral procesing regions, notably early and association visual and auditory regions. Such results are in line with findings in rats, where the electrical stimulation of the stomach elicits BOLD responses not only in somatosensory, insula, cingulate cortices, but also in motor, auditory and visual cortices (Cao et al., 2019). Changes in visual cortices activity has also been observed in humans in response to painful gastric distension (Van Oudenhove et al., 2009), subliminal rectal distension (Kern & Shaker, 2002), as well in dogs after gastric electrical stimulation (Yu et al., 2014). Similarly, colonic pain induces a response in the rat auditory cortex (Wang et al., 2008). The parabrachial nuclei, the major relay of both spinal and vagal visceral afferents, projects not only to thalamic relay nuclei involved in interoception and touch (Coen et al., 2012; Saper & Loewy, 1980), but also to the visual lateral geniculate nucleus (Erişir et al., 1997), and potentially to the auditory medial geniculate nuclei (Uhlrich et al., 1988). Projections of the parabrachial nucleus might thus at least partially mediate gastric-BOLD coupling in sensory cortices. It is also worth underlining that we find no gastric-BOLD coupling in anterior insular and medial prefrontal regions, two regions involved in parasympathetic control of the stomach in rats (Levinthal & Strick, 2020).

Beyond direct ascending or descending communication with the stomach, gastric-coupling could also stem from intra-cortical connectivity. For instance, direct connections have been reported between secondary somatosensory cortex to auditory regions, between visual area MT and primary somatosensory (Cappe & Barone, 2005). Finally, neuromodulation might also be involved (Rebollo et al., 2021; Rinaman & Schwartz, 2004). In particular, activity in the locus coeruleus, the main source of norepinephrine to the forebrain, is modulated by gastric, colonic and bladder distension (Elam et al., 1986; Saito et al., 2002), inducing fluctuations in arousal that can also be obtained with distension of the small intestine (Kukorelli & Juhász, 1977).

### Intersubject variability

We examined which individual variables, including anxiety often associated with cardiac interoception (Domschke et al., 2010), covary with gastric-brain coupling in the group of young, healthy participants studied here. We found either robust evidence for no covariation using Bayesian statistics, or no evidence for covariation, despite having a sample size large enough to detect moderate effect sizes. Our experimental setting was designed to minimize variation in digestive and autonomic state. Therefore, we cannot rule out the possibility that BMI, hunger or anxiety influence gastric-brain coupling in participants with larger BMIs or anxiety levels, or if we had explicitly manipulated anxiety or hunger in a within subject-design.

The only variable displaying a significant association with gastric-brain coupling was the LF-HRV/HF-HRV ratio. This ratio relates a mixture of both sympathetic and parasympathetic activity (LF-HRV) to a purely parasympathetic measure (HF-HRV). Thus, large values of this ratio might reflect either an increase in sympathetic activity, or an overall increase in both sympathetic and parasympathetic tone (Billman, 2013; Quintana et al., 2016). Both components could be involved here. Since somatomotor sympathetic cortical regions have the largest effect sizes of gastric-brain coupling, a sympathetic drive modulating both the LF-HRV/HF-HRV ratio and gastric brain coupling is possible. Additionally, our experimental set-up, with participants being neither hungry nor full, could involve a reduced parasympathetic tone.

### Possible functional roles of gastric-brain coupling

The precise function of the coupling between the BOLD signal and the gastric rhythm in this extended sensory and motor network remains highly speculative at this stage. Indeed, it takes a lot of information to define the function of a given brain region (Genon et al., 2018), let alone a whole network. Still, the spatial layout of the gastric network and the involvement of all sensory and motor cortices is puzzling, and calls for further interpretation. In the following, we consider several non-exclusive working hypotheses regarding the potential functional consequences of gastric-brain coupling.

The gastric network could be functionally related to interoception and autonomic processes. Somatosensory, motor, premotor, cingulate motor and insular cortices, which belong to the gastric network, have established roles in interoception and autonomic processes (Critchley et al., 2004; Dum et al., 2016; Levinthal & Strick, 2020), and the strength of gastric-brain coupling is related to an index of cardiac activity. Given the physiological function of the stomach, a specific link with feeding behavior should be considered. A recent study found an negative correlation between weight loss and power at 0.05 Hz in gastric network regions across 90 individuals undergoing a weight reduction intervention (Levakov et al., 2020), suggesting a link between energy regulation and brain activity at gastric frequency. Still, the coupling between gastric rhythm and activity in early auditory and visual regions is not readily explained by a functional role of the gastric network limited to bodily regulations.

Another view, that might encompass a link with energy regulation, is that gastric-brain coupling is related to arousal, given the anatomical and functional links reviewed above, as well as to the observantion that the amplitude of parieto-occipital alpha rhythm is coupled to the gastric rhythm in humans (Richter et al., 2017). Whether gastric-brain coupling correspond to fluctuations of arousal within one gastric cycle, akin to the notion “pulsed arousal” associated with the cardiac cycle (Garfinkel & Critchley, 2016), or to longer episodes of high or low arousal spanning several gastric cycles, remains to be determined. However this view does not account for the specific layout of the gastric network. Indeed, why should gastric-related fluctuations of arousal be concentrated in sensory and motor regions and spare most of the transmodal regions typically associated with conceptual, abstract processing (Behrens et al., 2013)?

Finally, gastric-brain coupling might reflect an overall scaffolding mechanism contributing to the organization of large-scale neural activity, involved in the coordination between brain regions to bind information (Fries, 2005; Singer & Gray, 1995) expressed in different formats and coordinates (Azzalini et al., 2019). The sensory and motor regions of the gastric network contain topographical representations of the body surface, activated by touch, movement or visual perception of body parts (Amiez & Petrides, 2014; Orlov et al., 2010; Penfield & Boldrey, 1937), as well as of external space, which is represented in retinotopic coordinates in visual cortices and in tonotopic space in auditory cortices. The gastric rhythm, acting as a common input to all those regions, could facilitate the alignment and coordination of the different coordinate systems in which external information from the senses is expressed – in other words, act as a binding mechanism facilitating between-area communication (Azzalini et al., 2019) – with the stomach delivering different time stamps to different regions in a mechanism reminiscent of traveling waves (Wang, 2010). Such a facilitation of inter-areal communication between sensory and motor regions would be an interesting complement to the known top-down control from cognitive areas to sensory regions.

## Conclusion

The multiple cortical areas of the human brain have long been thought be organized in an ascending hierarchy (Felleman & Van Essen, 1991) composed of relatively independent modules corresponding to each sensory modality converging onto higher-order, transmodal regions (Mesulam, 1998). While this view has been refined (see e.g., (Markov et al., 2013; Young, 1992), it is still much present in the narrative of large scale brain organization, and fits with the classical parcellation in distinct resting-state networks for different modalities (Yeo et al., 2011). Our findings show that regions considered to be mostly independent are actually all tightly linked through delayed functional connectivity with the stomach. The monitoring of visceral inputs should thus be integrated into current views of the cortical hierarchy.

## Acknowledgements

This work was supported by the European Research Council (ERC) under the European Union’s Horizon 2020 research and innovation program (grant agreement No 670325) to C.T.-B., as well as by ANR-17-EURE-0017. I.R. was supported by a grant from DIM Cerveau et Pensée and Fondation Bettencourt-Schueller. We thank Benoît Béranger and Juliette Klamm for their help during data acquisition.

