## Supplemental Table I and II for "The sensory and motor components of the cortical hierarchy are coupled to the rhythm of the stomach during rest" (from another version of this preprint: version 1, 2021-05-27)

### Supplementary Material

Table SI : MNI coordinates of regions synchronized to the stomach at the group level and the corresponding Automatic Anatomical Labeling (AAL) name

| **Cluster #** | **AAL Area** | **N voxels** | **mm^3^** | **% area** | **T in peak** | **x** | **y** | **z** |
| --- | --- | --- | --- | --- | --- | --- | --- | --- |
| 1 | Cingulum Mid R | 9 | 27 | 2,40 | 3,45 | 6 | -37 | 34 |
| 1 | Cingulum Post L | 24 | 72 | 30,46 | 5,18 | 0 | -34 | 28 |
| 1 | Cingulum Post R | 15 | 45 | 26,31 | 4,40 | 3 | -43 | 19 |
| 1 | Calcarine L | 250 | 750 | 65,07 | 4,96 | 3 | -94 | 4 |
| 1 | Calcarine R | 126 | 378 | 39,79 | 4,76 | 6 | -94 | 4 |
| 1 | Cuneus L | 167 | 501 | 64,31 | 4,63 | -3 | -76 | 31 |
| 1 | Cuneus R | 147 | 441 | 60,67 | 5,64 | 6 | -79 | 31 |
| 1 | Lingual L | 158 | 474 | 44,32 | 4,76 | -3 | -64 | 1 |
| 1 | Lingual R | 162 | 486 | 41,39 | 4,99 | 18 | -64 | -11 |
| 1 | Occipital Sup L | 49 | 147 | 21,08 | 4,28 | -18 | -85 | 34 |
| 1 | Occipital Sup R | 110 | 330 | 45,75 | 4,75 | 24 | -82 | 28 |
| 1 | Occipital Mid L | 41 | 123 | 7,37 | 4,03 | -24 | -91 | 13 |
| 1 | Occipital Mid R | 32 | 96 | 8,96 | 4,22 | 30 | -82 | 19 |
| 1 | Occipital Inf L | 13 | 39 | 8,12 | 3,90 | -15 | -97 | -8 |
| 1 | Fusiform L | 35 | 105 | 8,90 | 5,39 | -24 | -49 | -14 |
| 1 | Fusiform R | 44 | 132 | 10,27 | 4,52 | 27 | -70 | -8 |
| 1 | Parietal Sup R | 7 | 21 | 1,85 | 4,43 | 12 | -82 | 52 |
| 1 | Precuneus L | 84 | 252 | 13,99 | 5,12 | -6 | -46 | 10 |
| 1 | Precuneus R | 110 | 330 | 19,80 | 4,86 | 3 | -76 | 49 |
| 1 | Cerebelum Crus1 L | 129 | 387 | 29,12 | 5,25 | -27 | -76 | -26 |
| 1 | Cerebelum Crus2 L | 59 | 177 | 18,31 | 4,99 | -21 | -79 | -44 |
| 1 | Cerebelum 4 5 L | 27 | 81 | 14,10 | 4,10 | -15 | -55 | -14 |
| 1 | Cerebelum 4 5 R | 16 | 48 | 10,92 | 4,58 | 12 | -61 | -14 |
| 1 | Cerebelum 6 L | 48 | 144 | 16,65 | 4,38 | -24 | -67 | -29 |
| 1 | Cerebelum 6 R | 79 | 237 | 25,86 | 6,72 | 18 | -64 | -17 |
| 1 | Vermis 4 5 | 50 | 150 | 44,19 | 4,59 | 3 | -52 | 4 |
| 1 | Vermis 6 | 11 | 33 | 17,42 | 3,89 | 0 | -73 | -11 |
| 1 | Vermis 8 | 10 | 30 | 24,18 | 4,25 | 6 | -61 | -29 |
| 1 | Vermis 9 | 13 | 39 | 43,91 | 4,76 | 0 | -55 | -35 |
| 2 | Precentral R | 167 | 501 | 29,03 | 5,52 | 36 | -25 | 49 |
| 2 | Frontal Sup R | 17 | 51 | 2,46 | 4,64 | 15 | -16 | 73 |
| 2 | Supp Motor Area L | 91 | 273 | 24,91 | 4,58 | -6 | -7 | 61 |
| 2 | Supp Motor Area R | 97 | 291 | 24,04 | 4,42 | 3 | 5 | 70 |
| 2 | Cingulum Mid L | 31 | 93 | 9,39 | 3,83 | -3 | -7 | 46 |
| 2 | Cingulum Mid R | 33 | 99 | 8,80 | 4,00 | 9 | -22 | 46 |
| 2 | Postcentral R | 80 | 240 | 12,30 | 4,21 | 48 | -22 | 55 |
| 2 | Parietal Sup R | 24 | 72 | 6,35 | 4,08 | 21 | -55 | 70 |
| 2 | Paracentral Lobule L | 20 | 60 | 8,71 | 5,01 | 0 | -25 | 58 |
| 2 | Paracentral Lobule R | 12 | 36 | 8,44 | 4,29 | 15 | -28 | 58 |
| 3 | Precentral L | 155 | 465 | 25,83 | 5,68 | -24 | -13 | 73 |
| 3 | Frontal Sup L | 31 | 93 | 5,06 | 4,63 | -27 | -4 | 64 |
| 3 | Supp Motor Area L | 15 | 45 | 4,11 | 3,91 | -3 | -10 | 76 |
| 3 | Postcentral L | 106 | 318 | 16,01 | 4,37 | -21 | -34 | 61 |
| 3 | Paracentral Lobule L | 35 | 105 | 15,25 | 4,34 | -6 | -19 | 79 |
| 4 | Rolandic Oper R | 46 | 138 | 20,31 | 4,63 | 51 | -13 | 10 |
| 4 | Insula R | 52 | 156 | 17,26 | 4,54 | 36 | -7 | 10 |
| 4 | Postcentral R | 10 | 30 | 1,54 | 3,54 | 69 | -13 | 13 |
| 4 | SupraMarginal R | 14 | 42 | 4,17 | 3,70 | 48 | -34 | 25 |
| 4 | Heschl R | 24 | 72 | 56,64 | 5,37 | 51 | -13 | 7 |
| 4 | Temporal Sup R | 116 | 348 | 21,70 | 4,86 | 54 | -46 | 16 |
| 5 | Rolandic Oper L | 65 | 195 | 38,58 | 5,23 | -51 | -16 | 13 |
| 5 | Postcentral L | 58 | 174 | 8,76 | 4,88 | -63 | -22 | 22 |
| 5 | SupraMarginal L | 19 | 57 | 8,89 | 4,41 | -60 | -22 | 19 |
| 5 | Heschl L | 16 | 48 | 41,79 | 4,12 | -51 | -13 | 7 |
| 5 | Temporal Sup L | 70 | 210 | 17,92 | 4,99 | -60 | -19 | 10 |
| 6 | Occipital Inf R | 65 | 195 | 38,62 | 5,26 | 30 | -97 | -14 |
| 6 | Temporal Mid R | 8 | 24 | 1,07 | 3,76 | 54 | -73 | -2 |
| 6 | Temporal Inf R | 27 | 81 | 4,46 | 3,86 | 45 | -70 | -11 |
| 6 | Cerebelum Crus1 R | 42 | 126 | 9,32 | 4,70 | 51 | -73 | -26 |
| 7 | Fusiform R | 37 | 111 | 8,64 | 5,29 | 39 | -40 | -26 |
| 7 | Cerebelum Crus1 R | 17 | 51 | 3,77 | 4,40 | 36 | -55 | -35 |
| 7 | Cerebelum Crus2 R | 10 | 30 | 2,78 | 5,00 | 42 | -55 | -41 |
| 7 | Cerebelum 6 R | 45 | 135 | 14,73 | 4,76 | 33 | -55 | -35 |
| 8 | Precentral R | 74 | 222 | 12,86 | 4,91 | 45 | -10 | 49 |
| 8 | Frontal Mid R | 32 | 96 | 3,68 | 4,57 | 42 | 5 | 40 |
| 8 | Postcentral R | 22 | 66 | 3,38 | 4,39 | 60 | -22 | 49 |
| 8 | SupraMarginal R | 6 | 18 | 1,79 | 3,39 | 63 | -22 | 46 |
| 9 | Frontal Inf Oper R | 7 | 21 | 2,94 | 4,37 | 63 | 11 | 4 |
| 9 | Frontal Inf Orb R | 25 | 75 | 8,61 | 3,99 | 51 | 23 | -11 |
| 9 | Rolandic Oper R | 31 | 93 | 13,69 | 5,22 | 66 | -1 | 7 |
| 9 | Insula R | 8 | 24 | 2,66 | 3,50 | 51 | 8 | -5 |
| 9 | Temporal Sup R | 7 | 21 | 1,31 | 4,89 | 66 | 2 | 4 |
| 9 | Temporal Pole Sup R | 33 | 99 | 14,49 | 5,00 | 63 | 5 | 1 |
| 10 | Cingulum Mid L | 28 | 84 | 8,48 | 3,92 | 0 | -37 | 46 |
| 10 | Cingulum Mid R | 15 | 45 | 4,00 | 4,54 | 3 | -40 | 46 |
| 10 | Precuneus L | 19 | 57 | 3,16 | 4,38 | -6 | -43 | 61 |
| 10 | Paracentral Lobule L | 17 | 51 | 7,41 | 4,18 | -9 | -34 | 49 |
| 11 | Parietal Sup L | 15 | 45 | 4,27 | 3,79 | -18 | -64 | 58 |
| 11 | Precuneus L | 40 | 120 | 6,66 | 4,83 | -3 | -70 | 61 |
| 11 | Precuneus R | 13 | 39 | 2,34 | 3,73 | 3 | -70 | 61 |
| 12 | Precuneus L | 14 | 42 | 2,33 | 3,98 | 0 | -55 | 52 |
| 12 | Precuneus R | 46 | 138 | 8,28 | 4,36 | 9 | -46 | 43 |
| 12 | Paracentral Lobule R | 11 | 33 | 7,73 | 4,31 | 3 | -37 | 67 |
| 13 | Precentral L | 8 | 24 | 1,33 | 4,60 | -45 | -7 | 31 |
| 13 | Postcentral L | 61 | 183 | 9,21 | 5,41 | -54 | -13 | 34 |
| 14 | Frontal Sup Medial L | 28 | 84 | 5,50 | 3,85 | -9 | 59 | 7 |
| 14 | Frontal Sup Medial R | 16 | 48 | 4,41 | 4,52 | 6 | 62 | 1 |
| 14 | Frontal Med Orb L | 13 | 39 | 10,63 | 3,96 | -3 | 50 | -8 |
| 14 | Frontal Med Orb R | 8 | 24 | 5,49 | 3,58 | 6 | 59 | -2 |
| 15 | Temporal Mid R | 63 | 189 | 8,40 | 5,14 | 51 | -61 | 13 |
| 16 | Cerebelum Crus1 L | 44 | 132 | 9,93 | 5,23 | -42 | -73 | -35 |
| 16 | Cerebelum Crus2 L | 23 | 69 | 7,14 | 3,97 | -48 | -67 | -44 |
| 17 | Caudate L | 14 | 42 | 8,55 | 3,98 | -6 | 5 | 10 |
| 18 | Thalamus L | 39 | 117 | 20,84 | 4,45 | -6 | -19 | 7 |
| 18 | Thalamus R | 15 | 45 | 8,34 | 3,79 | 6 | -13 | 16 |
| 19 | Fusiform L | 12 | 36 | 3,05 | 3,64 | -30 | -58 | -17 |
| 19 | Cerebelum Crus1 L | 12 | 36 | 2,71 | 4,02 | -39 | -55 | -35 |
| 19 | Cerebelum 6 L | 42 | 126 | 14,57 | 4,63 | -36 | -58 | -26 |
| 20 | Postcentral R | 43 | 129 | 6,61 | 6,16 | 60 | -7 | 25 |
| 21 | Frontal Sup R | 41 | 123 | 5,94 | 5,45 | 21 | 32 | 55 |
| 22 | Cerebelum Crus1 R | 16 | 48 | 3,55 | 3,78 | 12 | -88 | -26 |
| 22 | Cerebelum Crus2 R | 25 | 75 | 6,94 | 4,34 | 15 | -85 | -32 |
| 24 | Supp Motor Area L | 24 | 72 | 6,57 | 3,94 | -6 | 14 | 61 |
| 24 | Supp Motor Area R | 26 | 78 | 6,44 | 4,92 | 6 | 14 | 46 |
| 24 | Cingulum Mid R | 6 | 18 | 1,60 | 3,82 | 9 | 11 | 43 |
| 25 | Temporal Sup L | 16 | 48 | 4,10 | 4,30 | -57 | 2 | -11 |
| 25 | Temporal Mid L | 27 | 81 | 3,21 | 5,38 | -57 | -7 | -11 |
| 26 | Postcentral L | 8 | 24 | 1,21 | 3,76 | -18 | -40 | 73 |
| 26 | Paracentral Lobule L | 33 | 99 | 14,38 | 5,44 | -6 | -34 | 73 |
| 27 | Caudate L | 20 | 60 | 12,22 | 4,22 | -9 | 11 | -8 |
| 27 | Putamen L | 19 | 57 | 11,07 | 5,11 | -21 | 11 | 4 |
| 28 | Frontal Inf Oper R | 8 | 24 | 3,36 | 3,78 | 48 | 14 | 19 |
| 28 | Frontal Inf Tri R | 25 | 75 | 6,83 | 4,44 | 51 | 23 | 25 |
| 29 | Rolandic Oper L | 9 | 27 | 5,34 | 4,21 | -39 | -13 | 19 |
| 29 | Insula L | 17 | 51 | 5,38 | 4,47 | -36 | -19 | 16 |
| 30 | Temporal Sup R | 33 | 99 | 6,17 | 4,97 | 63 | -7 | -5 |
| 32 | Frontal Sup R | 26 | 78 | 3,77 | 4,51 | 27 | -10 | 58 |

Table SII: Resting state networks according to the 7-network parcellation of Yeo et al (Yeo et al., 2011) and cortical regions synchronized with the stomach according to (Glasser et al., 2016) parcellation

| **RSN** | **Region name in atlas** | **Full name** | **OverlapLH** | **Effect sizes left** | **OverlapRH** | **Effect size right** |
| --- | --- | --- | --- | --- | --- | --- |
| Sensorimotor | 4 | Primary Motor Cortex | 47.8 | 0.82 | 47.7 | 0.78 |
| Sensorimotor | 3b | Primary Sensory Cortex | 50.4 | 0.74 | 52.9 | 0.79 |
| Sensorimotor | A1 | Primary Auditory Cortex | 61.8 | 0.67 | 64.8 | 0.61 |
| Sensorimotor | 5m | Area 5m | 62.0 | 0.53 | 7.0 | 0.46 |
| Sensorimotor | 5mv | Area 5 medio vental | 15.8 | 0.48 | 0.0 | - |
| Sensorimotor | 24dd | Dorsal Area 24d | 30.7 | 0.64 | 71.5 | 0.67 |
| Sensorimotor | 24dv | Ventral Area 24d | 5.8 | 0.37 | 25.7 | 0.54 |
| Sensorimotor | 1 | Area 1 | 53.2 | 0.77 | 38.2 | 0.75 |
| Sensorimotor | 2 | Area 2 | 22.6 | 0.65 | 1.0 | 0.39 |
| Sensorimotor | 3a | Area 3a | 45.5 | 0.70 | 57.6 | 0.77 |
| Sensorimotor | 6d | Dorsal area 6 | 59.4 | 0.75 | 62.7 | 0.71 |
| Sensorimotor | 6mp | Area 6mp | 27.9 | 0.56 | 24.8 | 0.59 |
| Sensorimotor | 6v | Ventral Area 6 | 9.9 | 0.49 | 15.1 | 0.45 |
| Sensorimotor | 43 | Area 43 | 79.0 | 0.68 | 37.8 | 0.61 |
| Sensorimotor | OP4 | Area OP4/PV | 64.2 | 0.61 | 35.3 | 0.50 |
| Sensorimotor | OP1 | Area OP1/SII | 73.6 | 0.61 | 25.9 | 0.56 |
| Sensorimotor | OP2-3 | Area OP2-3/VS | 29.3 | 0.55 | 41.2 | 0.54 |
| Sensorimotor | 52 | Area 52 | 25.6 | 0.52 | 64.2 | 0.62 |
| Sensorimotor | RI | RetroInsular Cortex | 80.4 | 0.65 | 24.9 | 0.54 |
| Sensorimotor | PFcm | Area PFcm | 57.6 | 0.58 | 40.8 | 0.51 |
| Sensorimotor | TA2 | Area TA2 | 0.8 | 0.40 | 47.8 | 0.66 |
| Sensorimotor | FOP2 | Frontal OPercular Area 2 | 1.3 | 0.34 | 29.8 | 0.54 |
| Sensorimotor | PBelt | ParaBelt Complex | 61.2 | 0.59 | 74.5 | 0.65 |
| Sensorimotor | A5 | Auditory 5 Complex | 19.0 | 0.58 | 36.5 | 0.79 |
| Sensorimotor | Ig | Insular Granular Complex | 42.3 | 0.57 | 83.0 | 0.63 |
| Sensorimotor | MBelt | Medial Belt Complex | 38.5 | 0.57 | 83.9 | 0.73 |
| Sensorimotor | LBelt | Lateral Belt Complex | 49.9 | 0.49 | 43.1 | 0.59 |
| Sensorimotor | A4 | Auditory 4 Complex | 20.7 | 0.50 | 47.3 | 0.63 |
| Visual | V1 | Primary Visual Cortex | 69.1 | 0.60 | 53.8 | 0.56 |
| Visual | V6 | Sixth Visual Area | 64.6 | 0.55 | 59.2 | 0.55 |
| Visual | V2 | Second Visual Area | 51.2 | 0.61 | 56.1 | 0.60 |
| Visual | V3 | Third Visual Area | 49.1 | 0.59 | 49.4 | 0.64 |
| Visual | V4 | Fourth Visual Area | 32.2 | 0.50 | 29.5 | 0.64 |
| Visual | V8 | Eighth Visual Area | 20.9 | 0.45 | 80.1 | 0.59 |
| Visual | V3A | Area V3A | 66.4 | 0.50 | 95.6 | 0.55 |
| Visual | V7 | Seventh Visual Area | 57.7 | 0.47 | 89.9 | 0.57 |
| Visual | FFC | Fusiform Face Complex | 0.0 | 0.25 | 23.0 | 0.72 |
| Visual | V3B | Area V3B | 80.1 | 0.47 | 13.5 | 0.43 |
| Visual | LO2 | Area Lateral Occipital 2 | 0.0 | - | 21.8 | 0.40 |
| Visual | PIT | Posterior InferoTemporal | 0.0 | - | 51.1 | 0.53 |
| Visual | MT | Middle Temporal Area | 0.0 | - | 26.7 | 0.48 |
| Visual | PreS | PreSubiculum | 0.0 | - | 6.9 | 0.42 |
| Visual | ProS | ProStriate Area | 69.0 | 0.60 | 44.9 | 0.54 |
| Visual | DVT | Dorsal Transitional Visual Area | 25.6 | 0.53 | 79.1 | 0.60 |
| Visual | V6A | Area V6A | 44.4 | 0.43 | 96.7 | 0.60 |
| Visual | VMV1 | VentroMedial Visual Area 1 | 30.0 | 0.46 | 50.0 | 0.57 |
| Visual | VMV3 | VentroMedial Visual Area 3 | 30.0 | 0.50 | 80.1 | 0.59 |
| Visual | V3CD | Area V3CD | 8.8 | 0.31 | 0.0 | - |
| Visual | LO3 | Area Lateral Occipital 3 | 0.0 | - | 17.0 | 0.38 |
| Visual | VMV2 | VentroMedial Visual Area 2 | 28.0 | 0.37 | 46.5 | 0.59 |
| Visual | VVC | Ventral Visual Complex | 25.2 | 0.67 | 31.9 | 0.64 |
| DMN | RSC | RetroSplenial Complex | 56.0 | 0.75 | 63.2 | 0.66 |
| DMN | SFROI | Superior Frontal Language Area | 26.2 | 0.50 | 24.7 | 0.51 |
| DMN | 7m | Area 7m | 0.0 | 0.30 | 13.0 | 0.46 |
| DMN | POS1 | Parieto-Occipital Sulcus Area 1 | 55.4 | 0.59 | 47.5 | 0.52 |
| DMN | 23d | Area 23d | 13.7 | 0.37 | 34.1 | 0.43 |
| DMN | v23ab | Area ventral 23 a+b | 55.1 | 0.51 | 37.4 | 0.53 |
| DMN | d23ab | Area dorsal 23 a+b | 38.6 | 0.47 | 51.9 | 0.49 |
| DMN | 31pv | Area 31p ventral | 0.6 | 0.24 | 8.0 | 0.35 |
| DMN | p32 | Area p32 | 0.0 | 0.33 | 7.7 | 0.44 |
| DMN | 10r | Area 10r | 17.8 | 0.44 | 11.5 | 0.47 |
| DMN | 8BROI | Area 8B Lateral | 0.0 | - | 16.7 | 0.53 |
| DMN | 10d | Area 10d | 21.2 | 0.57 | 36.8 | 0.62 |
| DMN | STGa | Area STGa | 0.0 | - | 12.0 | 0.57 |
| DMN | STSda | Area STSd anterior | 41.1 | 0.79 | 8.8 | 0.61 |
| DMN | STSdp | Area STSd posterior | 0.0 | - | 30.1 | 0.60 |
| DMN | STSvp | Area STSv posterior | 0.0 | - | 12.5 | 0.49 |
| DMN | TE1a | Area TE1 anterior | 4.9 | 0.32 | 0.3 | 0.34 |
| DMN | PGi | Area PGi | 0.0 | - | 7.4 | 0.42 |
| DMN | 31pd | Area 31pd | 0.0 | - | 15.9 | 0.42 |
| DMN | STSva | Area STSv anterior | 16.0 | 0.47 | 15.5 | 0.47 |
| DMN | TE1m | Area TE1 Middle | 0.0 | - | 23.5 | 0.54 |
| Saliency | 55b | Area 55b | 18.8 | 0.50 | 58.7 | 0.63 |
| Saliency | PSL | PeriSylvian Language Area | 0.5 | 0.33 | 6.6 | 0.49 |
| Saliency | STV | Superior Temporal Visual Area | 0.0 | - | 19.9 | 0.55 |
| Saliency | 5l | Area 5l | 7.4 | 0.52 | 20.0 | 0.55 |
| Saliency | 23c | Area 23c | 2.6 | 0.29 | 32.2 | 0.54 |
| Saliency | SCEF | Supplementary and Cingulate Eye Field | 53.8 | 0.70 | 73.3 | 0.68 |
| Saliency | 6ma | Area 6m anterior | 12.3 | 0.45 | 21.3 | 0.51 |
| Saliency | p24pr | Area Posterior 24 prime | 9.5 | 0.36 | 38.8 | 0.48 |
| Saliency | p32pr | Area p32 prime | 1.1 | 0.33 | 21.5 | 0.49 |
| Saliency | 6r | Rostral Area 6 | 0.5 | 0.56 | 22.9 | 0.57 |
| Saliency | PoI2 | Posterior Insular Area 2 | 0.5 | 0.41 | 36.4 | 0.67 |
| Saliency | FOP4 | Frontal OPercular Area 4 | 0.0 | - | 14.2 | 0.55 |
| Saliency | MI | Middle Insular Area | 0.0 | - | 21.1 | 0.50 |
| Saliency | FOP1 | Frontal OPercular Area 1 | 1.0 | 0.43 | 81.5 | 0.62 |
| Saliency | FOP3 | Frontal OPercular Area 3 | 0.0 | - | 11.8 | 0.37 |
| Saliency | TPOJ1 | Area TemporoParietoOcci pital Junction 1 | 0.0 | 0.32 | 7.0 | 0.52 |
| Saliency | PFop | Area PF opercular | 30.5 | 0.48 | 2.2 | 0.33 |
| Saliency | PoI1 | Area Posterior Insular 1 | 0.0 | - | 15.5 | 0.57 |
| Control | POS2 | Parieto-Occipital Sulcus Area 2 | 5.4 | 0.34 | 79.4 | 0.57 |
| Control | 7Pm | Medial Area 7P | 11.4 | 0.37 | 18.8 | 0.35 |
| Control | 8BM | Area 8BM | 2.0 | 0.20 | 10.4 | 0.49 |
| Control | 44 | Area 44 | 0.0 | - | 12.9 | 0.50 |
| Control | IFJa | Area IFJa | 0.0 | - | 23.4 | 0.42 |
| Control | IFSp | Area IFSp | 0.0 | - | 47.8 | 0.48 |
| Control | p9-46v | Area posterior 9-46v | 0.0 | - | 9.1 | 0.37 |
| Control | a9-46v | Area anterior 9-46v | 44.8 | 0.60 | 0.0 | - |
| Control | 9-46d | Area 9-46d | 6.7 | 0.53 | 0.0 | - |
| Control | a10p | Area anterior 10p | 43.1 | 0.74 | 0.0 | - |
| Control | 11ROI | Area 11l | 8.3 | 0.55 | 0.0 | - |
| Control | i6-8 | Inferior 6-8 Transitional Area | 2.6 | 0.25 | 9.5 | 0.40 |
| Control | s6-8 | Superior 6-8 Transitional Area | 0.0 | - | 23.0 | 0.53 |
| Control | TE1p | Area TE1 posterior | 0.0 | - | 10.4 | 0.47 |
| Control | 31a | Area 31a | 2.3 | 0.38 | 44.7 | 0.52 |
| Control | p10p | Area posterior 10p | 7.8 | 0.45 | 0.0 | - |
| Attention | FEF | Frontal Eye Fields | 29.8 | 0.60 | 52.7 | 0.62 |
| Attention | PEF | Premotor Eye Field | 0.0 | - | 11.0 | 0.44 |
| Attention | IPS1 | IntraParietal Sulcus Area 1 | 0.8 | 0.29 | 25.8 | 0.55 |
| Attention | PCV | PreCuneus Visual Area | 16.4 | 0.45 | 57.1 | 0.55 |
| Attention | 7Am | Medial Area 7A | 26.5 | 0.48 | 38.2 | 0.42 |
| Attention | 7PROI | Lateral Area 7P | 12.2 | 0.37 | 5.8 | 0.17 |
| Attention | 7PC | Area 7PC | 11.0 | 0.32 | 0.0 | - |
| Attention | VIP | Ventral IntraParietal Complex | 39.2 | 0.49 | 0.0 | - |
| Attention | 6a | Area 6 anterior | 13.3 | 0.52 | 0.3 | 0.46 |
| Attention | PFt | Area PFt | 1.7 | 0.34 | 3.8 | 0.29 |
| Attention | TPOJ3 | Area TemporoParietoOcci pital Junction 3 | 0.0 | - | 36.1 | 0.56 |
| Attention | PGp | Area PGp | 0.0 | - | 17.5 | 0.48 |
| Attention | IP0 | Area IntraParietal 0 | 12.2 | 0.34 | 2.4 | 0.42 |
| Limbic | 10v | Area 10v | 4.2 | 0.35 | 6.0 | 0.40 |
| Limbic | 10pp | Polar 10p | 30.2 | 0.60 | 0.0 | 0.37 |
